# Plasmodesmata-localized proteins and reactive oxygen species orchestrate light-induced rapid systemic signaling in Arabidopsis

**DOI:** 10.1101/2020.10.07.329995

**Authors:** Yosef Fichman, Ronald J. Myers, DeAna G. Grant, Ron Mittler

## Abstract

Systemic signaling and systemic acquired acclimation (SAA) are key to the survival of plants during episodes of abiotic stress. These processes depend on a continuous chain of cell-to-cell signaling events that extends from the initial tissue that senses the stress (local tissue) to the entire plant (systemic tissues). Among the different systemic signaling molecules and processes thought to be involved in this cell-to-cell signaling mechanism are reactive oxygen species (ROS), calcium, electric and hydraulic signals. How these different signals and processes are interlinked, and how they transmit the systemic signal all the way from the local tissue to the entire plant, remain however largely unknown. Here, studying the systemic response of *Arabidopsis thaliana* to a local treatment of excess light stress, we report that respiratory burst oxidase homolog D (RBOHD)-generated ROS enhance cell-to-cell transport and plasmodesmata (PD) pore size in a process that depends on the function of PD-localized proteins (PDLPs) 1 and 5, promoting the cell-to-cell transport of systemic signals during responses to light stress. We further identify aquaporins, and several different calcium-permeable channels, belonging to the glutamate receptor-like, mechanosensitive small conductance-like, and cyclic nucleotide-gated families, as involved in this process, but determine that their function is primarily required for the maintenance of the signal in each cell along the path of the systemic signal, as well as for the establishment of acclimation at the local and systemic tissues. PD and RBOHD-generated ROS orchestrate therefore light stress-induced rapid cell-to-cell spread of systemic signals in Arabidopsis.

**One-sentence summary:** Respiratory burst oxidase homolog D-generated reactive oxygen species enhance cell-to-cell transport and plasmodesmata (PD) pore size in a process that depends on the function of the PD-localized proteins (PDLPs) 1 and 5, promoting the cell-to-cell transport of rapid systemic signals during the response of Arabidopsis to excess light stress.

## Introduction

Acclimation of plants to changes in their environment requires many different physiological, molecular and metabolic responses. These are controlled by multiple signal transduction cascades, hormonal signaling pathways, and changes in the steady-state level of calcium and reactive oxygen species (ROS, *1–4*). In addition to activating acclimation mechanisms at the specific plant tissue(s) exposed to stress, different abiotic stresses, as well as mechanical injury, can trigger rapid systemic signaling pathways that result in systemic acquired acclimation (SAA), or systemic wound responses (SWR), at the whole-plant level (*5–18*). In the case of abiotic stresses, such as excess light or heat stresses, the process of rapid systemic signaling and SAA result in the protection of plants from a subsequent exposure to abiotic stress (*9, 14, 19–22*).

Among the many different systemic signaling pathways thought to mediate rapid systemic responses and SAA or SWR are electrical, calcium, hydraulic, and ROS waves (*6–8, 10, 12, 13, 21–23*). Electrical signals and calcium waves were recently shown to be dependent on the function of the calcium-permeable glutamate receptor-like (GLR) GLR3.3 and GLR3.6 channels during SWR (*7, 10, 13, 23*), and calcium and ROS waves were proposed to be linked through the function of the respiratory burst oxidase homolog D (RBOHD) protein during systemic responses to light and salt stress (*6, 15, 17, 24, 25*). The RBOHD protein was further shown to be required for the propagation of different electric signals in response to light stress (*14*). Although it is unknown how hydraulic waves are linked to electric, calcium and ROS waves, it was proposed that calcium (and/or other ion/cation)-permeable mechanosensitive channel of small conductance-like (MSL, *26*) channels could sense hydraulic waves at the systemic tissues and convert them into calcium signals (*4, 17, 24*). Calcium signals can further impact ROS signals via the function of many different calcium-binding proteins and/or calcium-dependent kinases/phosphatase switches, or by directly binding to the EF-binding domains of the RBOHD protein (*1–4, 24, 25*). Other calcium-permeable channels proposed to be involved in regulating local and/or systemic responses to stress include cyclic nucleotide-gated ion channels (CNGCs, *27*), annexins (ANN, *28*), reduced hyperosmolality-induced [Ca^2+^]_i_ increase (OSCA, *29*), and two-pore channel (TPC, *30*).

Activation of RBOHD results in the generation of superoxide (O_2_^−^) molecules that dismutate into hydrogen peroxide (H_2_O_2_) at the apoplast, and aquaporins such as plasma membrane (PM) intrinsic protein channels (PIPs, *31, 32*) were demonstrated to regulate H_2_O_2_ translocation across the PM from the apoplast into the cytosol. The translocation of H_2_O_2_ into the cytosol enables it to alter different redox-dependent reactions, kinases/phosphatase molecular switches, and/or calcium-permeable channels, further driving different local and systemic acclimation pathways (*6, 8, 9, 19–22, 25, 31*). This type of apoplast-to-cytosol translocation of signaling molecules such as H_2_O_2_ is also responsible for cell-to-cell communication of systemic signals during SAA (*6, 8, 9, 18–22*). A signaling compound such as H_2_O_2_, produced by one cell could therefore enter a neighboring cell and trigger acclimation and defense mechanisms in it, resulting in the activation of its own RBOHD, and a chain of cell-to-cell transmission of this type of signal (*i.e*., the ROS wave) could mediate rapid systemic signaling (*6, 8, 14, 19–22, 33–35*). In addition to such an apoplastic route of H_2_O_2_–to-cytosol translocation among different cells during systemic signaling, a symplastic plasmodesmata (PD)-dependent route of systemic signaling could also be playing a role in the translocation of systemic signals, mediating different calcium, redox or kinases/phosphatase switch modes between cells (*7, 17, 36–38*). Proteins such as PD-localized proteins 1 and 5 (PDLP1, PDLP5, *39*), PM-localized leucine-rich-repeat receptor-like-kinase 7 (KIN7, *40*) and GFP-arrested trafficking 1 (GAT1, *41*) were for example shown to regulate translocation through PDs. However, the role of these proteins in regulating systemic responses at the rapid rate required for SAA to be effective (*4*) was not demonstrated. In addition, although PD pore size was proposed to be regulated by changes in redox levels (*42–45*), whether or not PD transport is regulated by ROS during rapid systemic signaling and SAA to excess light stress is currently unknown. Here, using a newly developed method to image ROS signals in live plants grown in soil (*8, 20–22, 46*), coupled with different grafting and acclimation studies, we studied the regulation of systemic signals in wild type and many different *Arabidopsis thaliana* mutants impaired in ROS, calcium, PD and aquaporin functions (table S1) in response to a local application of light stress.

## Results

### The role of calcium-permeable channels in regulating rapid systemic ROS signals in Arabidopsis

Calcium and ROS signaling are thought to co-regulate many of the different responses of plants to changes in environmental conditions (*1, 2, 4*). The double mutant *glr3.3glr3.6* was previously reported to be impaired in the propagation of systemic electric signals and Ca^2+^ waves in response to wounding (*7, 10, 13, 23*), highlighting GLR3.3GLR3.6 as an important hub for systemic signaling. Interestingly, in our hands, and in response to a local excess light stress treatment (not injury, as in *7, 10, 13, 23*), the *glr3.3glr3.6* double mutant was not deficient in rapid systemic ROS signaling, and only displayed a suppressed systemic ROS wave response to a local application of high light (HL) stress (Fig. 1A and movie S1). Moreover, the single *glr3.3* mutant displayed an enhanced rate of systemic ROS signal propagation, whereas the single *glr3.6* mutant was similar to wild-type, in response to the local application of HL (fig. S1 and table S1). In contrast to the double *glr3.3glr3.6* mutant, two independent alleles each of *cngc2* that could potentially regulate RBOHD (Fig. 1B and movie S2), or *msl2* that could potentially link hydraulic waves with calcium and ROS signals (Fig. 1C and movie S3), were completely deficient in the induction and/or propagation of the rapid systemic ROS signal in response to a local HL treatment. Although similar results were found with two independent alleles each of *msl3* (fig. S2), two independent alleles each of a large number of other mutants for Ca^2+^-permeable channels, including *msl10, ann1, osca1*, and *tpc1*, displayed an enhanced rate of systemic ROS signal propagation in response to a local application of HL (figs. S3 to S6 and table S1). These results suggest that many different Ca^2+^-permeable channels are involved in regulating the formation and/or propagation of systemic ROS signals in plants. Some, such as *glr3.3glr3.6, msl2, msl3*, and *cngc2*, are required, whereas others, for example, *ann1, tpc-1, msl10* and *osca1*, could play a repressive or inhibitory role. The findings that at least 3 different types of Ca^2+^-permeable channels (GLR, CNGC and MSL, Fig. 1) are required for mediating rapid systemic ROS signaling in Arabidopsis underscores the tight level of regulation and coordination required for this process to occur.

**Fig. 1.**
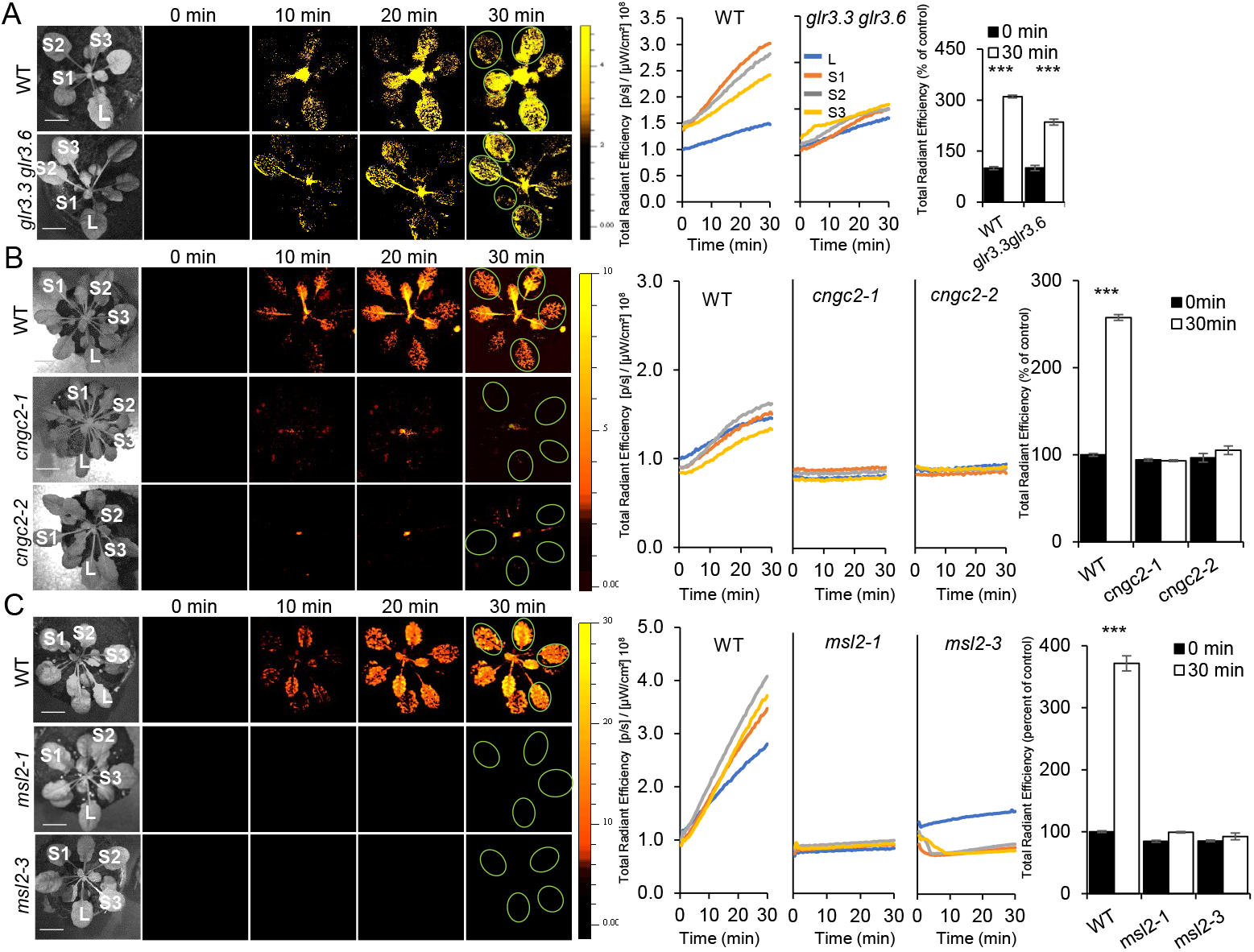
Three different types of Ca^2+^-permeable channels are required for light stress-induced rapid systemic ROS signaling in Arabidopsis. (**A**) Representative time-lapse images of systemic ROS accumulation in wild-type and *glr3.3glr3.6* double mutant *Arabidopsis thaliana* plants subjected to a 2 min local (L) high light (HL) stress treatment (applied to leaf L only), are shown on left, representative line graphs showing continuous measurements of ROS levels in local and systemic (S) leaves over the entire course of the experiment (0 to 30 min) are shown in the middle (ROIs used for calculating them are indicated with light green circles on the images to the left), and statistical analysis of ROS accumulation in local and systemic leaves of all plants used for the analysis at 0 and 30 min is shown on right. (**B**) Same as in (A), but for wild-type and two independent alleles of *cngc2*. (**C**) Same as in (A), but for wild-type and two independent alleles of *msl2*. All experiments were repeated at least 3 times with 10 plants per biological repeat. Student t-test, SE, N=12, ***P < 0.005. Scale bar indicates 1 cm. *Abbreviations used*: CNGC, cyclic nucleotide-gated ion channel, GLR, glutamate receptor-like, HL, high light, L, local, MSL, mechanosensitive channel of small conductance-like, ROI, region of interest, S, systemic.

### The role of PD and aquaporins in regulating rapid systemic ROS signals in Arabidopsis

Although much is known about calcium and ROS integration during responses to changes in abiotic and biotic conditions (*1–4, 6, 7, 10, 13, 15, 24*), less is known about the role of PD and aquaporins in these responses. To determine whether HL-induced systemic signaling in Arabidopsis utilizes an aquaporin-associated apoplastic, or a PD-dependent symplastic, route for its initiation and/or propagation, we measured HL-induced systemic ROS signals in different mutants impaired in aquaporin or PD functions. Mutants for *pdlp1* or *pdlp5* (two independent alleles of each), were impaired in mediating the rapid systemic ROS signal in response to a local application of HL (Figs. 2A, 2B, movies S4, S5 and table S1). In contrast, two other PD mutants, *kin7* and *gat1* (two independent alleles of each), displayed an enhanced, or wild-type-like, rates of systemic ROS signal propagation, respectively (figs. S7, S8 and table S1). Two independent alleles of the aquaporin mutant *pip1;2* had an enhanced rate of systemic ROS signal propagation (Fig. 2C, movie S6 and table S1), while two independent alleles of the aquaporin *pip2;1* were completely deficient in the initiation or propagation of the rapid systemic ROS signal (Fig. 2D, movie S7 and table S1). In contrast, *pip1;4* (two independent alleles) displayed a wild-type-like systemic ROS response (fig. S9 and table S1). The finding that PIP2;1, that localizes to vascular bundles of Arabidopsis (*31, 32*), is essential for the initiation and/or propagation of the ROS wave (Fig. 2D, movie S7 and table 1), is in agreement with our recent findings that rapid systemic ROS signaling occurs via phloem and xylem parenchyma cells during systemic responses to excess light stress (*22*). Similar to calcium-permeable channels (Fig. 1), the function of different aquaporins (that could mediate ROS movement between the apoplast and the cytosol, *31, 32*), and PD-related proteins (that could regulate the movement of redox state, Ca^2+^ and other signals between cells, *37–45*), might be required within the same cell, or within different cells, for mediating rapid systemic signals in plants. Our findings indicate therefore that both symplastic and apoplastic routes could be required for rapid systemic signaling in Arabidopsis.

**Fig. 2.**
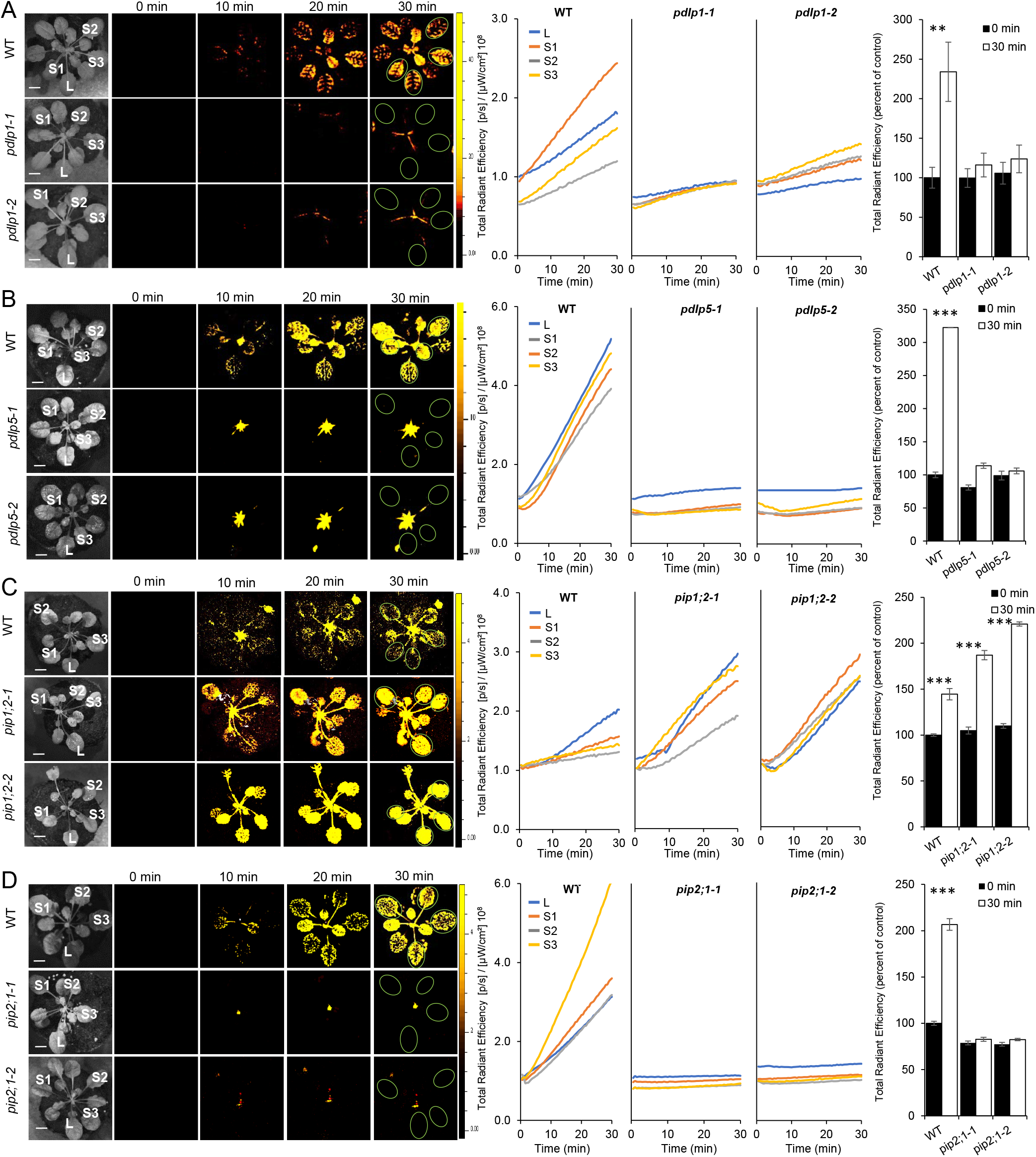
The PD-localized proteins PDLP1 and PDLP5 and the aquaporin protein PIP2;1 are required for light stress-induced rapid systemic ROS signaling in Arabidopsis. (**A**) Representative time-lapse images of systemic ROS accumulation in wild-type and *pdlp1* (two independent alleles) *Arabidopsis thaliana* plants subjected to a 2 min local (L) high light (HL) stress treatment (applied to leaf L only), are shown on left, representative line graphs showing continuous measurements of ROS levels in local and systemic (S) leaves over the entire course of the experiment (0 to 30 min) are shown in the middle (ROIs used for calculating them are indicated with light green circles on the images to the left), and statistical analysis of ROS accumulation in local and systemic leaves of all plants used for the analysis at 0 and 30 min is shown on right. (**B**) Same as in (A), but for wild-type and two independent alleles of *pdlp5*. (**C**) Same as in (A), but for wild-type and two independent alleles of *pip1;2*. (**D**) Same as in (A), but for wild-type and two independent alleles of *pip2;1*. All experiments were repeated at least 3 times with 10 plants per biological repeat. Student t-test, mean ± SE, N=12, ***P < 0.005, **P < 0.01. Scale bar indicates 1 cm. *Abbreviations used*: HL, high light, L, local, PDLP, plasmodesmata localized protein, PIP, plasma membrane intrinsic protein, ROI, region of interest, S, systemic.

### SAA of mutants deficient in calcium-permeable channels, PD and aquaporin functions

The triggering of SAA by systemic signals in plants is thought to play a key role in plant survival during episodes of abiotic stress (*1–4, 9, 14, 19–22*) To determine whether the block or suppression in rapid systemic signaling displayed in the different mutants described above (Figs. 1 and 2) affected acclimation to excess light stress (*9, 14, 19–22*), we tested the systemic and local acclimation of *glr3.3glr3.6, cngc2, msl2, pip2;1* and *pdlp5* to HL stress, following a local treatment of HL stress. All mutants tested were impaired, albeit at various levels, in systemic or local acclimation to excess light stress (Fig. 3A). Analysis of transcript expression in local and systemic tissues of these mutants, in response to a local HL stress treatment, revealed that the steady-state level of different acclimation transcripts such as *ZAT10* and *ZAT12* (*6, 9, 20, 21*) was enhanced in local leaves (that were directly subjected to the HL treatment) of all mutants. With the exception of *glr3.3glr3.6*, that displayed suppressed expression of acclimation transcripts in systemic leaves, the expression of *ZAT10* and *ZAT12* in systemic tissues of all other mutants was repressed (Fig. 3B). In addition, and in agreement with the lack of systemic tissue acclimation to excess light stress (Fig. 3A), the expression of *MYB30*, that is required for systemic, but not local, acclimation to excess light stress (*21*), was repressed in the systemic tissues of all mutants (Fig. 3B). This finding demonstrates that the different mutants tested are able to sense the stress at their local tissues, exposed to stress, but are unable to transmit (or have a suppressed transmission rate, in the case of *glr3.3glr3.6*), the systemic signal from their local stressed tissues to their systemic leaves. The disruption or suppression in systemic signaling, caused by mutations in the *GLR3.3GLR3.6, CNGC2, MSL2, PIP2;1* or *PDLP5* genes (Figs. 1, 2 and table S1), was therefore detrimental for plant acclimation to stress (Fig. 3A), underscoring the important biological role these proteins play in this process.

**Fig. 3.**
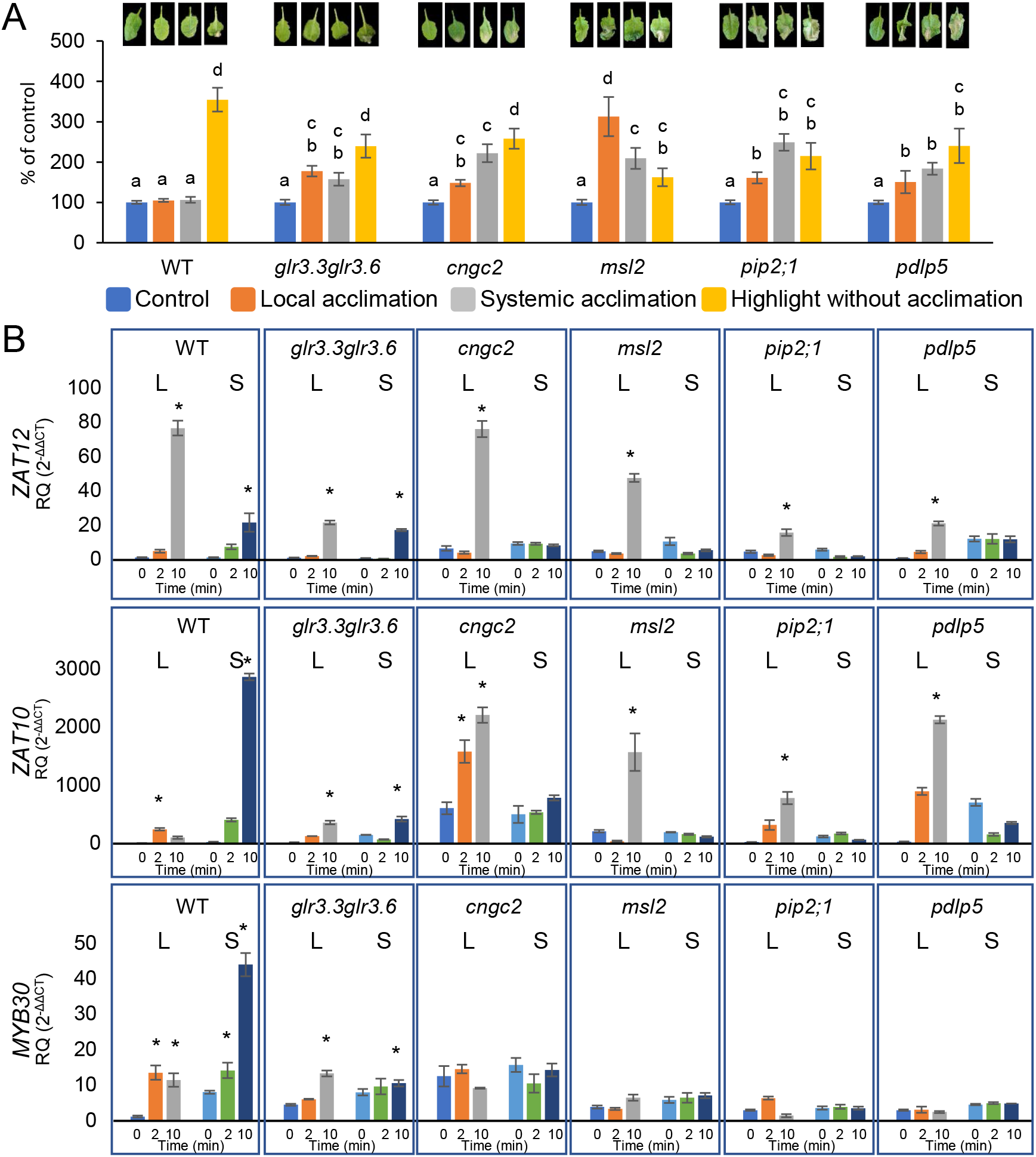
Acclimation of mutants impaired in systemic ROS signaling to light stress. (**A**) Representative leaf images (top) and measurements of leaf injury (increase in ion leakage, bottom) for wild-type and the *glr3.3glr3.6, cngc2, msl2,pip2;1* and *pdlp5* mutants. Measurements are shown for unstressed plants (control), local leaves subjected to a pretreatment of high light (HL) stress, followed by a long HL stress period (local acclimation), systemic leaves of plants subjected to a pretreatment of HL applied to their local leaves, followed by a long HL stress period (systemic acclimation), and systemic leaves of plants subjected to a long HL stress period without pretreatment (highlight without acclimation). Results are presented as percent of control (leaves not exposed to the light stress treatment). (**B**) Real-time quantitative PCR analysis of transcript expression in local and systemic leaves of wild-type and the *glr3.3glr3.6, cngc2, msl2,pip2;1* and *pdlp5* mutants subjected to a local 0, 2 or 10 min HL treatment. Transcripts tested (*ZAT12, ZAT10, MYB30*) were previously found to respond to HL stress at the local and systemic leaves of wild-type plants (*6, 9, 20, 21*). Results, expressed in RQ (relative quantity), were obtained by normalizing relative transcript expression and comparing it to control wild type from local leaf. All experiments were repeated at least 3 times with at least 10 plants per biological repeat. Acclimation experiments were analyzed using a one-way ANOVA followed by a Tukey’s post hoc test, and transcript expression analysis was analyzed using a student t-test, mean ± S.E. *Abbreviations used*: CNGC, cyclic nucleotide-gated ion channel, EL, electrolyte leakage, GLR, glutamate receptor-like, HL, high light, L, local, MSL, mechanosensitive channel of small conductance-like, PCR, polymerase chain reaction, PDLP, plasmodesmata localized protein, PIP, plasma membrane intrinsic protein, RQ, relative quantity, S, systemic.

### Determining the role of the different calcium-permeable channels, PD and aquaporin proteins in systemic signaling using a grafting approach

Because the suppressed ability of mutants impaired in *GLR3.3GLR3.6, CNGC2, MSL2, PIP2;1* or *PDLP5* function to systemically acclimate to HL (Fig. 3) could result from their inability to initiate and/or propagate the rapid systemic signal, we conducted grafting experiments (*47*) between wild-type plants and the *glr3.3glr3.6, cngc2, msl2,pip2;1,pdlp5* and *pdpl1* mutants. As controls we conducted grafting experiments between wild-type and wild-type plants, or wild-type and the *rbohD* mutant, that has reduced systemic apoplastic ROS accumulation in response to a local HL stress (*6, 8*), and is unable to induce SAA to HL (*9, 18*) or transmit a heat stress-induced systemic signal generated in a wild-type stock into its mutant scion (*14*). Only grafting events that remained green and maintained turgor level for up to 10 or more days, an indication of successful grafting (*47*), were studied, and multiple successful grafting events were obtained for all mutants described above. As shown in Fig. 4 and fig. S10, the *rbohD* mutant was unable to mediate the systemic ROS signal in the scion or the graft indicating that it is absolutely required for initiating and propagating the systemic ROS signal. The systemic ROS signal did not therefore initiate at the *rbohD* stock, nor did it propagate through the *rbohD* scion, following a local HL treatment of the *rbohD* or wild-type stocks, respectively. The only other mutants that displayed a similar behavior were *pdlp1* and *pdlp5*, indicating that PD function is also absolutely required along the entire path of the systemic signal (Fig. 4 and fig. S10). In contrast, the grafting results for the *glr3.3glr3.6, cngc2, msl2* and *pip2;1* mutants suggest that these proteins may have a cell-autonomous function (Fig. 4 and fig. S10). They may not be required for the initiation or the propagation of the systemic ROS signal, since the signal can be initiated and propagated though a stock of these mutants (that do not show enhanced ROS accumulation), and proceed through this stock into a wild-type scion in which it will cause systemic ROS accumulation (Fig. 4 and fig. S10). In contrast, however, once the systemic ROS signal is initiated in a wild-type stock, it will not further propagate though a scion made from these mutants and these scion sections would not display enhanced systemic ROS accumulation (Fig. 4 and fig. S10). *RBOHD, PDLP5*, and *PDLP1* are therefore required for the initiation and propagation of the rapid systemic ROS signal between cells, whereas *GLR3.3GLR3.6, CNGC2, MSL2* and *PIP2;1* are required for the amplification and/or maintenance of the systemic ROS signal in each individual cell (Fig. 4 and fig. S10).

**Fig. 4.**
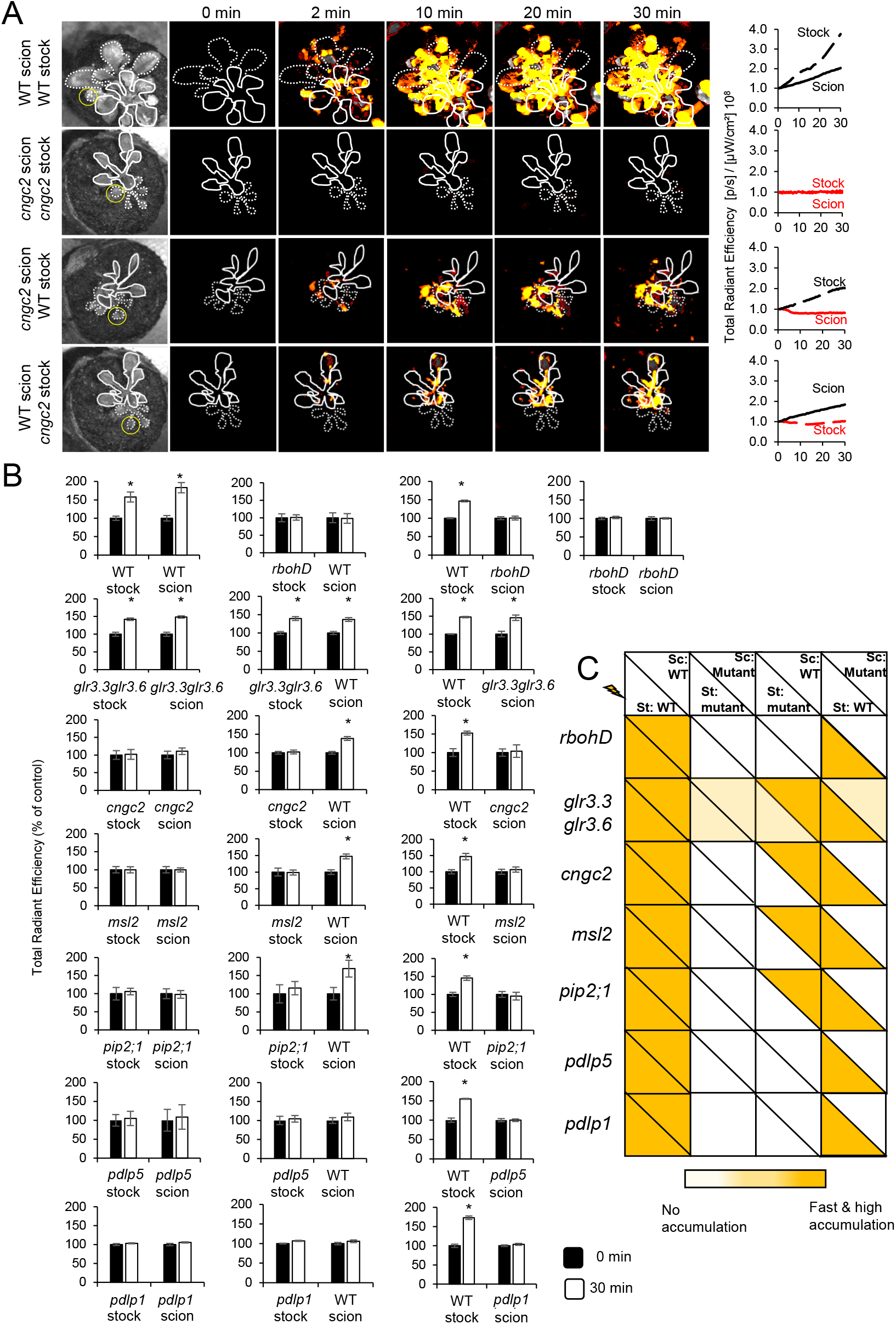
The PD-localized proteins PDLP1 and PDLP5, and RBOHD, are required for the initiation and propagation of the rapid systemic ROS signal. (**A**) Representative time-lapse images of systemic ROS accumulation in different grafting combinations between wild-type and the *cngc2* mutant, along with line graphs showing continuous measurements of ROS levels in the stock and the scion of these grafting combination (ROIs used for calculating them are indicated with light green circles on the images to the left), are shown as an example (examples of line graphs for all other grafting combinations are shown in fig. S10). The areas of local stock leaves subjected to a 2 min light stress are indicated by yellow circles, scions are indicated by solid white lines, and stocks are indicated by dashed white lines. (**B**) Statistical analysis of ROS signal intensity in the stock and scion of all plants used for the different grating combinations between wild-type and mutants impaired in systemic signaling (*rbohD, glr3.3glr3.6, cngc2, msl2, pip2;1, pdlp1* and *pdlp5*), 30 min following a 2 min high light stress treatment of a single stock leaf (local leaf). (**C**) Heat map summarizing the results obtained for the different grafting experiments. Color intensity indicates the presence and level of a ROS signal in the stock or scion parts of each grafting combination tested. All experiments were repeated at least 3 times with 10 plants per biological repeat (mean ± S.E., *p < 0.05, Student t-test). *Abbreviations used*: CNGC, cyclic nucleotide-gated ion channel, GLR, glutamate receptor-like, MSL, mechanosensitive channel of small conductance-like, PDLP, plasmodesmata localized protein, PIP, plasma membrane intrinsic protein, RBOHD, respiratory oxidase burst homolog D, ROI, region of interest, ROS, reactive oxygen species, Sc, scion, St. Stock, WT, wild-type.

### *RBOHD* and *PDLP5* impact cell-to-cell spread of carboxyfluorescein and PD pore area size during light stress in Arabidopsis

The findings that *RBOHD, PDLP1* and *PDLP5* are required for the propagation of the rapid systemic ROS signal from local to systemic tissues (Fig. 4 and fig. S10), could suggest that ROS and PD functions are interlinked during systemic signaling. Elevated ROS levels were recently proposed to enhance PD and tunneling nanotube transport in plant and mammalian cells (*45*). If such a mechanism occurs in Arabidopsis in response to excess light stress, it could explain why RBOHD and PDLPs are both required for systemic ROS signaling in Arabidopsis (Figs. 2 to 4). To test this possibility we measured the cell-to-cell spread of the fluorescent compound carboxyfluorescein (*48*) in local and systemic leaves of wild-type, *rbohD* and *pdlp5* plants in the presence or absence of a 2 min HL treatment applied to the local leaf. While carboxyfluorescein spread was facilitated in response to a local HL treatment in petioles and leaf cells of local and systemic leaves of wild-type plants, a similar response was not observed in the petioles and leaf cells of local or systemic leaves of *rbohD* and *pdlp5* plants (Figs. 5B and 5C). Because ROS were proposed to cause an enhancement in PD pore size, facilitating PD transport by a factor of ~10 (*45*), we used transmission electron microscopy (TEM, *49*) to measure PD pore area size in petioles of local leaves from wild-type, *rbohD* and *pdlp5* plants treated or untreated for 2 min with HL stress (focusing on vascular bundle and parenchyma cells). While the pore area size of PD (H/M- and X/Y-shaped) from wild-type plants increased in response to the HL stress treatment (in agreement with the facilitated spread of carboxyfluorescein following the excess light stress treatment; Figs. 5B and 5C), the PD pore area size of *rbohD* and *pdlp5* plants decreased (Fig. 5D, in agreement with the lack of HL-driven facilitated carboxyfluorescein spread in *rbohD* and *pdlp5* plants, Figs. 5B and 5C). The results presented in Fig. 5 suggest therefore that RBOHD-generated ROS impact cell-to-cell transport and PD pore area size in a PDLP5-dependent manner.

**Fig. 5.**
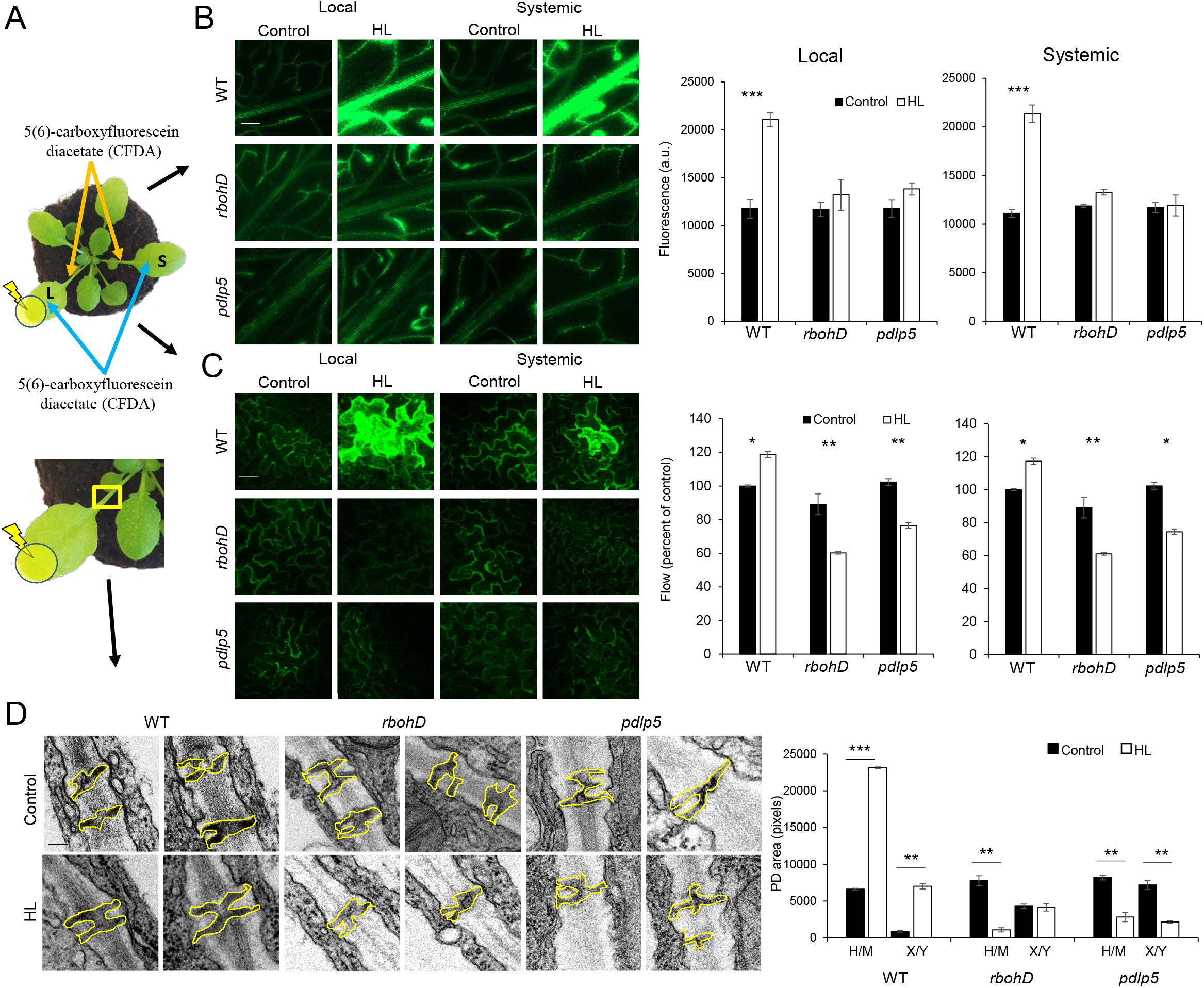
*RBOHD* and *PDLP5* impact cell-to-cell spread of carboxyfluorescein and PD pore area size during light stress responses in Arabidopsis. (**A**) The experimental designs used to measure carboxyfluorescein fluorescence spread through the vascular bundles (B) or cells (C) of local and systemic leaves of plants subjected to a HL stress treatment applied to leaf L only (Top), and PD pore size in petioles of leaves subjected to HL stress by transmission electron microscopy (TEM, D). (**B**) Representative images (left) and statistical analysis (right) of carboxyfluorescein fluorescence intensity in vascular bundles of wild-type (WT), *rbohD* and *pdlp5* petioles (local and systemic), subjected to a 2 min local light stress treatment. Plants were treated or untreated with light stress and incubated for 30 min. Petioles of detached leaves were then submerged in 5(6)-carboxyfluorescein diacetate (CDFA) for 5 min. Following incubation, carboxyfluorescein fluorescence intensity was measured at the vascular bundles of petioles 5 mm from the detachment sight. All experiments were repeated 6-10 times with 10 biological repeats. Student t-test, mean ± SE, N=96, ***P < 0.005. Scale bar indicates 250 μm. (**C**) Representative images (left) and statistical analysis (right) of carboxyfluorescein fluorescence flow between different cell layers in local and systemic leaf cells of wild-type (WT), *rbohD* and *pdlp5* plants, subjected to a 2 min local light stress treatment. Plants were treated or untreated with light stress and a drop of 5 μL CFDA was placed on the adaxial surfaces of the local and systemic leaves. Following 30 min of incubation, Z-scan of the epidermis tissues were obtained by confocal laser scanning microscope, and the rate of flow was calculated based on the increase in the number of fluorescent layers as the CFDA spread in the Z-axis between cells (*48*) comparing treated and untreated plants. All experiments were repeated 6 times with 10 biological repeats. Student t-test, mean ± SE, N=96, **P < 0.01, *P < 0.05. Scale bar indicates 100 μm. (**D**) TEM analysis of PD pore area in the petioles of local WT, *rbohD* and *pdlp5* leaves, subjected to a 2 min local light stress treatment (applied to the leaf area only). Representative PD images are shown on left and statistical analysis of PD pore area (H/M- and X/Y-shaped) is shown on right. All experiments were repeated 6-10 times with 10 biological repeats. Student t-test, mean ± SE, N=108, ***P < 0.005, **P < 0.01. Scale bar indicates 0.1 μm.

## Discussion

The findings presented in our study suggest that PD regulation (PDLP5, PDLP1) and ROS production (RBOHD) are required along the entire path of the HL stress-induced rapid systemic ROS signal (Figs. 2 to 5). In contrast, the GLR3.3GLR3.6 calcium-permeable channels may only have a supportive role in mediating HL-induced systemic ROS signals along the entire path (Figs. 1, 3 and 4). It is further proposed that the function of two different pathways is required to mediate the rapid systemic ROS signal from its initiation site to the entire plant and induce SAA to HL stress: *i*) a cell-autonomous pathway that amplifies the systemic ROS signal, triggers acclimation responses, and requires RBOHD, GLR3.3GLR3.6, PIP2;1, CNGC2, and MSL2, and *ii*) a cell-to-cell pathway that propagates the systemic signal and requires RBOHD, PDLP5, and PDLP1 (Fig. 6). Because mutants impaired in *GLR3.3GLR3.6, CNGC2, MSL2*, or *PIP2;1* were able to sense the HL stress at their local tissues, but failed to enhance the expression of different acclimation transcripts at their systemic leaves, as well as failed to induce acclimation in their local or systemic leaves (Fig. 3), and because stocks made from these mutants were able to transfer the systemic signal to a wild-type scion, but did not accumulate high ROS levels (Fig. 4 and fig. S10), it is possible that the role of the cell-autonomous pathway is to enhance the ROS signal, activate the expression of acclimation transcripts, and induce acclimation, in each cell along the path of the HL-induced systemic signal (Fig. 6). In contrast, PDLP5, PDLP1 and RBOHD, that are essential for transferring the systemic signal from the stock to the scion, as well as for local and systemic accumulation of ROS, the enhanced expression of acclimation transcripts in systemic tissues, and local and systemic acclimation (Figs. 2 to 5 and fig. S10), are required for both cell-to-cell systemic signal propagation and activation of acclimation responses during the systemic response of Arabidopsis to a local treatment of HL stress (Fig. 6). Because RBOHD function (ROS production) is required for both pathways (Fig. 6), it is likely that ROS propagate from cell-to-cell, or that ROS produced in each cell along the path of the systemic signal are required for propagating the systemic signal from cell-to-cell (Fig. 6). These functions could be mediated by low levels of ROS that are below the detection limit of the whole-plant ROS imaging method used. In contrast, the activation of acclimation responses within each cell along the path of the signal (cell-autonomous pathway) could require high levels of ROS that are detected by the ROS imaging method used (Fig. 4).

**Fig. 6.**
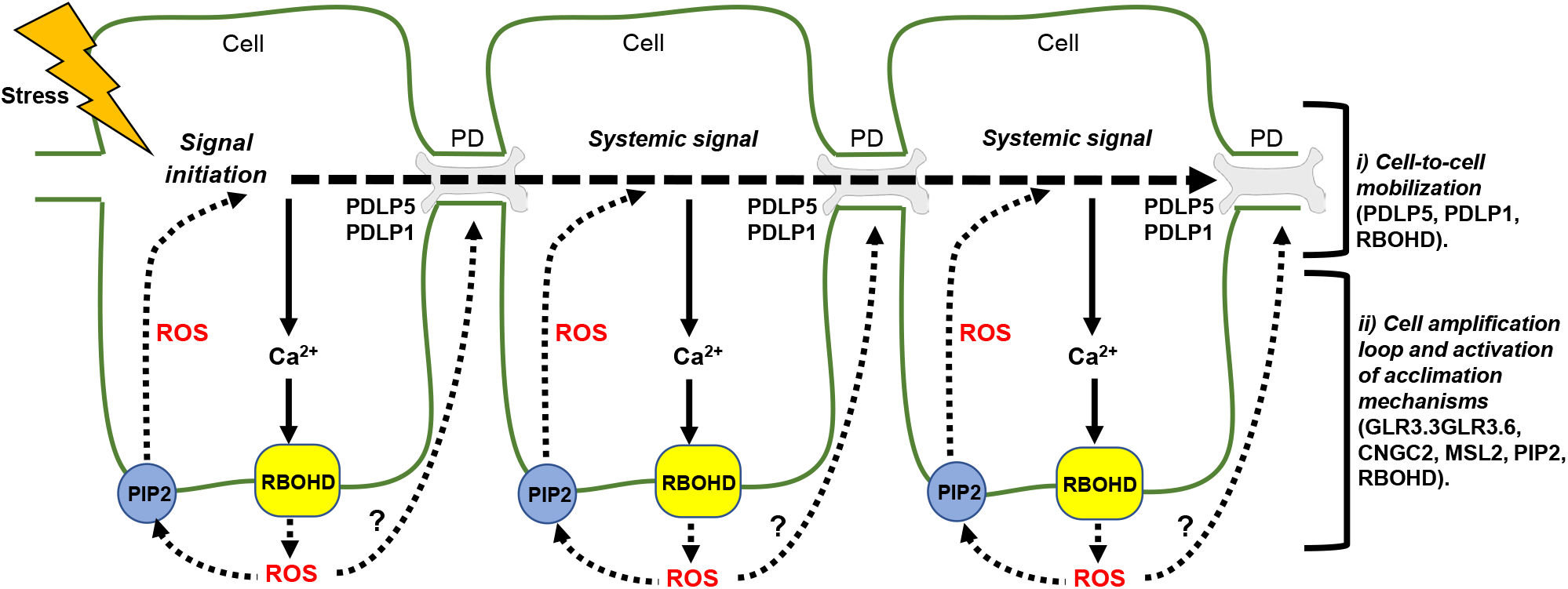
A hypothetical model for ROS and PD interactions during systemic signaling in Arabidopsis. A hypothetical model for the regulation of rapid systemic ROS signaling by the different Ca^2+^-permeable channels, ROS produced by RBOHD, PD and aquaporin functions. Two pathways are proposed to regulate rapid systemic ROS signaling in plants: *i*) a cell-to-cell pathway that involves PDLP5, PDLP1 and RBOHD, and *ii*) a cell-autonomous pathway (cell amplification loop) that amplifies and regulates the ROS and Ca^2+^ signals in each cell and involves RBOHD, GLR3.3GLR3.6, CNGC2, MSL2, and PIP2. ROS produced by each cell along the path of the signal are depicted as regulating PD function. More detailed discussion is provided in the text. *Abbreviations used*: CNGC, cyclic nucleotide-gated ion channel, GLR, glutamate receptor-like, MSL, mechanosensitive channel of small conductance-like, PD, plasmodesmata, PDLP, plasmodesmata localized protein, PIP, plasma membrane intrinsic protein, RBOHD, Respiratory burst oxidase homolog D, ROS, reactive oxygen species.

How could ROS be mobilized between cells, or affect the transport of the rapid systemic signal? RBOHD-produced ROS could accumulate at the apoplast and enter the cells that produced them, or neighboring cells, via PIP2;1, or they can be mobilized between cells via PD. Because PIP2;1 is not required for mobilizing the rapid systemic ROS signal though the stock to the scion in our grafting experiments, but PDLP5 and PDLP1 are (Fig. 4 and fig. S10), it is likely that ROS that enter cells back through PIP2;1 are mobilized between cells via PD (Fig. 6). Alternatively, ROS could affect PD function from their cytosolic or apoplastic side (*45*), and/or impact the oxidation state and function of PDLPs (*39, 42–44*), and enable rapid transport of the systemic signal (Fig. 6). ROS could of course use another, yet unidentified, route to enter neighboring cells. The recently proposed role for ROS in enhancing PD and tunneling nanotube transport in plant and mammalian cells (*45*), based in part on earlier studies in plants (*42–44*), could provide a key explanation to the role of ROS in cell-to-cell communication. The RBOHD-mediated production of ROS along the path of the systemic signal could play an important role in opening PD and promoting the transport of the systemic signal from cell-to-cell (Fig. 5). PDLPs were previously shown to be required for pathogen-induced PD closure responses that involve callose deposition (*37–39, 42–44*), in an apparent conflict with the results presented in our study (Figs. 2 to 5). Because these PD closure responses, as well as callose deposition, could take hours to days to develop (*37–39, 42–44*), while the responses reported in this study occur within minutes (Figs. 1 to 5), it is possible that PDLPs have multiple functions that may occur at different time-scales or rates. In response to a HL stress-induced short burst of ROS they are involved in PD opening (Fig. 5), while in response to prolonged ROS levels, and/or other pathogen-derived signals, they induce long-term PD closure (*37–39, 42–44*).

The findings that a deletion in a number of other calcium-permeable channels, PD and aquaporin proteins (*i.e., MSL10, ANN1, OSCA1, TPC1, KIN7*, and *PIP1;2*) results in enhanced propagation rates of systemic ROS signals, hints to the existence of additional, yet unknown, pathways that suppress systemic signals and/or alter their signatures. The remarkable differences observed between the spread of systemic ROS signals in the *pip1;2* and the *pip2;1* mutants (Figs. 2C and 2D) for example could result from differential interaction of these two channels with other proteins in the cell (*32, 50*), the differential stability of these two water channels in the presence of ROS (*51*), and/or the differential permeability of these two channels for H_2_O_2_ (*52*). In addition, because PIP2;1 primarily localizes to the vascular tissues of Arabidopsis (*31, 32*), it could be directly required for mediating the ROS wave process, that propagates through these same tissues (*22*). Systemic ROS signals could therefore be controlled by multiple different pathways, some promoting them and some suppressing or altering their signature (*3, 4, 8, 24*). Supporting the existence of ROS wave-suppressing pathways is a recent study showing that the transcription factor MYB30 regulates a pathway that suppresses systemic ROS signals in response to excess light stress (*21*).

Previous reports underscored the importance of the GLR3.3GLR3.6 calcium-permeable channels in mediating systemic wound responses (*7, 10, 13, 23*). In contrast, our findings reveal that these channels may only have a supportive role in mediating systemic signals in response to HL stress (Figs. 1, 3 and 4). This discrepancy raises an interesting possibility that different systemic signal transduction pathways are triggered by different stimuli. While some systemic signaling pathways, triggered for example by wounding, are absolutely dependent on GLR3.3GLR3.6 (*7, 10, 13, 23*), others, such as those triggered by excess light stress, do not (Figs. 1, 3 and 4). Interestingly, at least in our hands, both wounding- and HL-triggered systemic signal transduction pathways require ROS production by RBOHD (*8*). An alternative explanation is of course that during responses to HL stress the role of GLR3.3GLR3.6 is replaced by other calcium permeable channels such as CNGC2, or MSL2 (Figs. 1, 3 and 4).

Taken together, the findings presented in this work highlight a key role for PD and ROS in mediating systemic signals and SAA to excess light stress. In addition, they suggest that RBOH-generated ROS could enhance cell-to-cell transport and PD pore size in a process that depends on the function of PDLPs. Such a mechanism could control the mobilization of many different systemic signals in plants, triggered by different abiotic, biotic, or developmental cues.

## Materials and Methods

### Plant material, growth conditions and stress treatments

Homozygous *Arabidopsis thaliana* knockout lines (table S1) and wild-type plants were germinated and grown on peat pellets (Jiffy International, Kristiansand, Norway) under controlled conditions of 10hr/14hr light/dark regime, 50 μmol photons s^−1^m^−2^ and 21°C for 4 weeks. Plants were subjected to HL stress by illuminating a single leaf with 1700 μmol photons s^−1^m^−2^ using a ColdVision fiber optic LED light source (Schott, Southbridge, MA, USA), as described earlier (*8, 9, 18, 22*).

### Grafting

Adding a scion to a seedling stock was performed according to published literature (*47*). Briefly, seeds were germinated on 0.5X Murashige and Skoog (MS) media plates. An incision was made in seven-day-old stock seedlings to insert a scion into the cut while keeping the rosette of the stock plant intact. MS plates were incubated for five days in a growth chamber at 20°C under constant light (50 μmol photons s^−1^m^−2^; 20°C). Surviving grafted plants were transplanted to peat pellets and grown as described above for 5 more days before stress treatments. For each knockout line, four combinations were constructed and tested: wild-type (WT) as the scion and the stock, the mutant line as the scion and the stock, mutant scion on WT stock, and WT scion on a mutant stock. Grafting was repeated 40 times for each combination of each line with approximately 40% yield.

### ROS imaging

Plants were fumigated for 30 min with 50 μM 2’,7’-dichlorofluorescein diacetate (H2DCFDA, Millipore-Sigma, St. Louis, MO, USA) in a glass container using a nebulizer (Punasi Direct, Hong Kong, China) as previously reported (*8, 20–22*). Following fumigation, a local light stress treatment was applied to a single leaf for 2 min. Images of dichlorofluorescein (DCF) fluorescence were acquired using an IVIS Lumina 5 apparatus (PerkinElmer, Waltham, MA, USA) for 30 min. ROS accumulation was analyzed using Living Image 4.7.2 software (PerkinElmer) utilizing the math tools. Time course images were generated and radiant efficiency of regions of interest (ROI) were calculated. Radiant efficiency is defined as fluorescence emission radiance per incident excitation and is expressed as (p/s)/(μW/cm^2^), p-photons, sec-seconds, μW-micro-Watt, cm^2^-square centimeter (*8, 20–22*). In each experiment, wild type was compared as control to one set of mutants representing a gene of interest. Because the initial intensity of the ROS signal was sometimes different between different experiments, depending on the physiological state of plants, time of day, and the phenotype of the mutants, the visualization range scale was set for each experiment separately (*8, 20–22*). Visualization range scale was first set automatically by the computer based on the peak intensity of the entire experiment, and then corrected manually so that the progression rate will be visualized and not saturated (*8, 20–22*). This resulted in line graphs that were sometimes different in their initial start point between different experiments. All readings were therefore standardized to the same start point in all line graphs and all bar graphs are expressed as % of control (WT at 0 min). Each data set includes standard error of 8-12 technical repeats and a Student t-test score (*8, 20–22*). Dye penetration controls, shown in fig. S11, were performed by fumigation of plants with 0.3% hydrogen peroxide for 10 min following the H2DCFDA fumigation and acquisition of images in the IVIS Lumina S5 (*8, 20–22*).

### Systemic acquired acclimation assays

Leaf injury following light stress was measured using the electrolyte leakage assay, as described previously (*9, 14, 18, 20–22*). Briefly, systemic acclimation to HL stress was tested by exposing a local leaf to light stress for 10 min, incubating the plant under controlled conditions for 50 min and then exposing the same leaf (local) or another younger leaf (systemic) to HL stress (1700 μmol photons s^−1^m^−2^) for 45 min. Electrolyte leakage was measured by immersing the sampled leaf in distilled water for 1 hr and measuring water conductivity. Samples were then boiled, cooled down to room temperature and measured again for conductivity (total leakage). The electrolyte leakage was calculated as percentage of the conductivity before heating the samples over that of the boiled samples conductivity. Results are presented as percent of control (electrolyte leakage from leaves not exposed to the light stress treatment). Experiments consisted of 5 repeats for each condition in each line. Standard error was calculated using Microsoft Excel, one-way ANOVA (confidence interval = 0.05) and Tukey post hoc test were performed with IBM SPSS 25.

### Transcript expression

To measure the transcriptional response of local and systemic leaves to HL stress in 4-week-old plants, HL was applied to a single leaf for 2- or 10-min. Exposed leaf (local) and unexposed fully developed younger leaf (systemic) were collected for RNA extraction. RNA was extracted using Plant RNeasy kit (Qiagen, Hilden, Germany) according to the manufacture instructions. Quantified total RNA was used for cDNA synthesis (PrimeScript RT Reagent Kit, Takara Bio, Kusatsu, Japan). Transcript expression was quantified by real-time qPCR using iQ SYBR Green supermix (Bio-Rad Laboratories, Hercules, CA, USA), as described in (*6, 9, 14, 18, 21*), with specific primers for: *ZAT12* (AT5G59820) 5’-TGGGAAGAGAGTGGCTTGTTT-3’ and 5’-TAAACTGTTCTTCCAAGCTCCA-3’, *ZAT10* (AT1G27730) 5’-ACTAGCCACG TTAGCAGTAGC-3’ and 5’-GTTGAAGTTTGACCGGAAGTC-3’, and *MYB30* (AT3G28910) 5’-CCACTTGGCGAAAAAGGCTC-3’ and 5’-ACCCGCTAGCTGAGGAAGTA-3’ Elongation factor 1 alpha (5’-GAGCCCAAGTTTTTGAAGA-3’ and 5’-TAAACTGTTCTTCCAAGCTCCA-3’) was used for normalization of relative transcript levels. Results, expressed in RQ (relative quantity) were obtained by normalizing relative transcript expression and comparing it to control wild type from local leaf. The data represents 15 biological repeats and 3 technical repeats for each reaction. Standard error and Student t-test were calculated with Microsoft Excel.

### Spread of the fluorescent compound carboxyfluorescein among petiole and leaf cells following a local HL stress treatment

A PD permeability assay was carried out based on (*48*), using two different experimental settings. In the first experimental design, a single local leaf was subjected to HL stress for 2 min and plants were then incubated under normal conditions for 30 min to allow the systemic the signal to spread. The local leaf and a systemic leaf from the same plant were then cut, and their petioles dipped in 1 mM 5(6)-Carboxyfluorescein diacetate (CFDA, Ex/Em 492/517 nm, Millipore-Sigma, St. Louis, MO, USA) for 5 min. CFDA is a membrane-permeable dye which upon cell entry hydrolyzes to the fluorescent carboxyfluorescein compound. Leaves were then imaged using a Lionheart FX (BioTek, Winnoski, VT, USA) fluorescent microscope at X10 magnification using the GFP filter settings. Fluorescence intensity was measured at the vascular bundles of leaves 5 mm from the detachment sight using the Lionheart FX Gen5 image analysis mode (BioTek, Winnoski, VT, USA). Untreated plants were used as control with 6 biological repeats. In the second experimental design, following the local 2 min application of HL stress, a drop of 5 μL of CFDA was placed on the adaxial surfaces of the local and systemic leaves (*48*). Following 30 min of incubation, Z-scan of the epidermis tissues were obtained by confocal laser scanning microscope (Leica TCS SP8, Leica Microsystems, Wetzlar, Germany), and the rate of flow was calculated based on the increase in the number of fluorescent layers as the CFDA spread in the Z-axis between cells (*48*), comparing treated and untreated plants. Confocal images were acquired at the University of Missouri Molecular Cytology Core facility.

### Transmission electron microscopy (TEM)

Local leaves of four-week-old plants were subjected to a 2 min HL stress and processed for TEM as described in (*49*). Briefly, leaves were sampled and their petioles sectioned and fixed in 2% paraformaldehyde and 2% glutaraldehyde in 100 mm sodium cacodylate buffer, pH 7.35. Samples were incubated at 4°C for 1 hr, rinsed with cacodylate buffer followed by distilled water. En bloc staining was performed using 1% aqueous uranyl acetate at 4°C overnight. Samples were then rinsed with distilled water. A graded dehydration series was performed using ethanol, transitioned into acetone, and dehydrated tissues were then infiltrated with a 1v/1v of Epon and Spurr resin for 24 hr at room temperature and polymerized at 60°C overnight. Sections were cut to a thickness of 80 nm using an ultramicrotome (Ultracut UCT, Leica Microsystems, Wetzlar, Germany) and a diamond knife (Diatome, Hatfield, PA, USA). Images were acquired with a JEOL JEM 1400 transmission electron microscope (JEOL, Tokyo, Japan) at 80 kV on a Gatan Ultrascan 1000 CCD (Gatan, Inc., Pleasanton, CA, USA). The size of PDs in different vascular bundle and parenchyma cells was analyzed using ImageJ. Each experiment included 10 technical repeats and 20 biological repeats. Preparation of the samples and imaging were performed at the Electron Microscopy Core facility at the University of Missouri.

### Statistical analysis

Statistical analysis for ROS accumulation (total radiant efficiency), real-time quantitative PCR transcript expression, carboxyfluorescein fluorescence and PD pore area size measurements was performed by two-sided student t-test, and results are presented as mean ± SE, *p < 0.05, **p < 0.01, ***p < 0.001. Statistical analysis for acclimation studies was performed by a one-way ANOVA followed by a Tukey post hoc test, and results are presented as mean ± SE. Different letters denote statistical significance at p < 0.05.

## Supplementary Materials

Fig. S1. Imaging of the systemic ROS signal in the individual *glr3.3* or *glr3.6* mutants suggest that the GLR3.3 or GLR3.6 genes are not required for mediating light stress-induced systemic signaling in Arabidopsis.

Fig. S2. The calcium-permeable channel MSL3 is required for mediating light stress-induced systemic signaling in Arabidopsis.

Fig. S3. Enhanced systemic ROS signal in mutants impaired in the MSL10 calcium-permeable channel during light stress-induced systemic signaling in Arabidopsis.

Fig. S4. Enhanced systemic ROS signal in mutants impaired in the ANN1 calcium-permeable channel during light stress-induced systemic signaling in Arabidopsis.

Fig. S5. Enhanced systemic ROS signal in mutants impaired in the OSCA1 calcium-permeable channel during light stress-induced systemic signaling in Arabidopsis.

Fig. S6. Enhanced systemic ROS signal in mutants impaired in the TPC1 calcium-permeable channel during light stress-induced systemic signaling in Arabidopsis.

Fig. S7. Enhanced systemic ROS signal in mutants impaired in the plasmodesmata-regulating protein KIN7 during light stress-induced systemic signaling in Arabidopsis.

Fig. S8. Plasmodesmata-regulating protein GAT1 is not required for mediating light stress-induced systemic signaling in Arabidopsis.

Fig. S9. Aquaporin PIP1;4 is not required for mediating light stress-induced systemic signaling in Arabidopsis.

Fig. S10. Plasmodesmata function and apoplastic ROS production are required for both initiation and propagation of the systemic signal.

Fig. S11. Systemic ROS imaging following hydrogen peroxide fumigation in the tested plants (as control for dye penetration in the different mutants).

Table S1. Detailed description of the different alleles used in this study.

Movie S1. Time-lapse video imaging of systemic ROS accumulation in wild-type and *glr3.3glr3.6* plants in response to a 2 min local treatment of high light stress.

Movie S2. Time-lapse video imaging of systemic ROS accumulation in wild-type and *cngc2* plants in response to a 2 min local treatment of high light stress.

Movie S3. Time-lapse video imaging of systemic ROS accumulation in wild-type and *msl2* plants in response to a 2 min local treatment of high light stress.

Movie S4. Time-lapse video imaging of systemic ROS accumulation in wild-type and *pdlp1* plants in response to a 2 min local treatment of high light stress.

Movie S5. Time-lapse video imaging of systemic ROS accumulation in wild-type and *pdlp5* plants in response to a 2 min local treatment of high light stress.

Movie S6. Time-lapse video imaging of systemic ROS accumulation in wild-type and *pip1;2* plants in response to a 2 min local treatment of high light stress.

Movie S7. Time-lapse video imaging of systemic ROS accumulation in wild-type and pip2;1 plants in response to a 2 min local treatment of high light stress.

## Acknowledgments

We thank seeds donations by E. Farmer, C. Maurel, and G. Stacy. We thank the Arabidopsis Biological Resource Center for all additional seeds used in this study.

## Author contributions

Y.F. and R.J.M. performed experiments and analyzed the data. D.G.G. conducted TEM analysis. R.M. and Y.F. designed experiments, analyzed the data and wrote the manuscript.

## Data Availability

Data supporting the findings of this study is available from the corresponding author upon request.

