## Supplementary Figures and Table for "Plasmodesmata-localized proteins and reactive oxygen species orchestrate light-induced rapid systemic signaling in Arabidopsis"

**Table S1. Detailed description of the different alleles used in this study.**

| Genotype | Accession no. | Systemic response to highlight | mutant accessions | Subcellular localization | Expression |
|---|---|---|---|---|---|
| WT | | unchanged | | | |
| *rbohd* | AT5G47910 | inhibited | | PM | Leaf |
| *glr3.3_ glr3.6* | | suppressed | SALK_099757 X SALK_091801 | PM, M, ER, G, N X PM, M, ER, G, CH | Phloem X Xylem |
| *glr3.3* | AT1G42540 | enhanced | SALK_099757 | PM, M, ER, G, N | Phloem |
| *glr3*.6 | AT3G51480 | unchanged | SALK_091801 | PM, M, ER, G, CH | Xylem |
| *cngc2* | AT5G15410 | inhibited | SALK_019922C<br>SALK_066908C | PM, CH | Leaf, shoot |
| *msl2* | AT5G10490 | inhibited | CS69609<br>CS69611 | PM, M, CH, N | sperm cell, endosperm, embryo, leaf |
| *msl3* | AT1G58200 | inhibited | CS69719<br>SALK_201695C | M, C, N | Endosperm, leaf, mesophyll, root phloem, pollen |
| *msl10* | AT5G12080 | enhanced | SALK_076254<br>SAIL_292_A11 | PM, M, CH, N, P | Carpel, shoot, ovule, pistil, xylem |
| *pip1;2* | AT2G45960 | enhanced | SALK_019794C<br>SALK_0145347 | PM, M, CH, N, ER, G, P, V | Root endodermis, shoot vascular bundle sheath |
| *pip1;4* | AT4G00430 | unchanged | SAIL_75_F07<br>SAIL_1166_B06 | PM, M, C, CH, N, ER, G, P | Abscission zone, pistil, shoot apical meristem, flower, leaf |
| *pip2;1* | AT3G53420 | inhibited | *pip2;1-1* (AMAZE collection)<br>SM_3_35928 | PM, M, CH, N, ER, G, V | Root endodermis, petiole epidermis, shoot vascular bundle sheath, stem |
| *pdlp1* | AT5G43980 | inhibited | SAIL_515_B10<br>SM_3_36596 | PM, M, ER, G | Endosperm, leaf, cambium, mesophyll |
| *pdlp5* | AT1G70690 | inhibited | SALK_044770.1<br>SAIL_46_E06 | PM, CH, N, ER, G, V | Root xylem, leaf, stem |
| *kin7* | AT3G02880 | enhanced | SALK_019840C<br>GT_5_108995 | PM, M, CH, ER, G, V | Leaf, mesophyll, root |
| *gat1* | AT2G15570 | unchanged | SALK_078093<br>SAIL_793_B04.1 | M, C, N | Root, shoot apex, petal |

| | | | | | |
|---|---|---|---|---|---|
| *ann1* | AT1G35720 | enhanced | SALK_015426C<br>GABI_327B12 | PM, C, M, CH, P | Root, stem |
| *osca1* | AT4G04340 | enhanced | SALK_038633C<br>SAIL_523_G10 | PM, M, CH, ER, G | Sperm cell, leaf pavement cell, bundle sheath |
| *tpc1* | AT4G03560 | enhanced | SALK_074094<br>SALK_125650 | PM, M, N, ER, G | Root hair cell, leaf pavement cell, bundle sheath, petiole epidermis |

*Abbreviations used:* Apex, apical meristem; C, cytosol; CH, chloroplasts; ER, endoplasmic reticulum; G, Golgi; M, mitochondria; N, nucleus; P, peroxisome; PM, plasma membrane; V, vacuole.

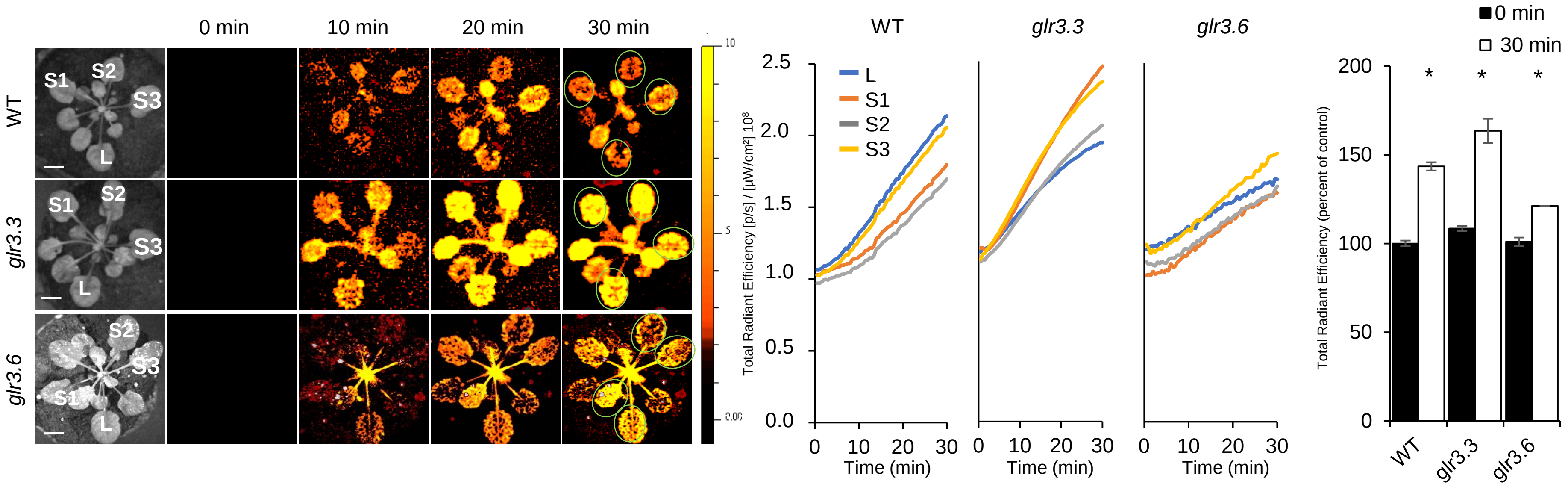


**Fig. S1. Imaging of the systemic ROS signal in the individual *glr3.3* or *glr3.6* mutants suggest that the GLR3.3 or GLR3.6 genes are not required for mediating light stress-induced systemic signaling in Arabidopsis.**

Representative time-lapse imaging of systemic ROS accumulation in wild-type, *glr3.3* and *glr3.6 Arabidopsis thaliana* plants subjected to a 2 min local (L) high light (HL) stress treatment (applied to leaf L only), is shown on left; representative line graphs of continuous measurements of ROS levels in the local (L) and systemic (S) leaves over the entire course of the experiment are shown in the middle (ROIs used for calculating them are indicated with light green circles on the images to the left); and statistical analysis of ROS accumulation in local and systemic leaves of all plants used for the analysis at 0 and 30 min is shown on right. All experiments were repeated at least 3 times with 10 plants per biological repeat. Student t-test, SE, N=12, *P < 0.05. Scale bar indicates 1 cm. *Abbreviations used:* HL, high light; L, local; GLR, glutamate receptor; ROI, region of interest; S, systemic.

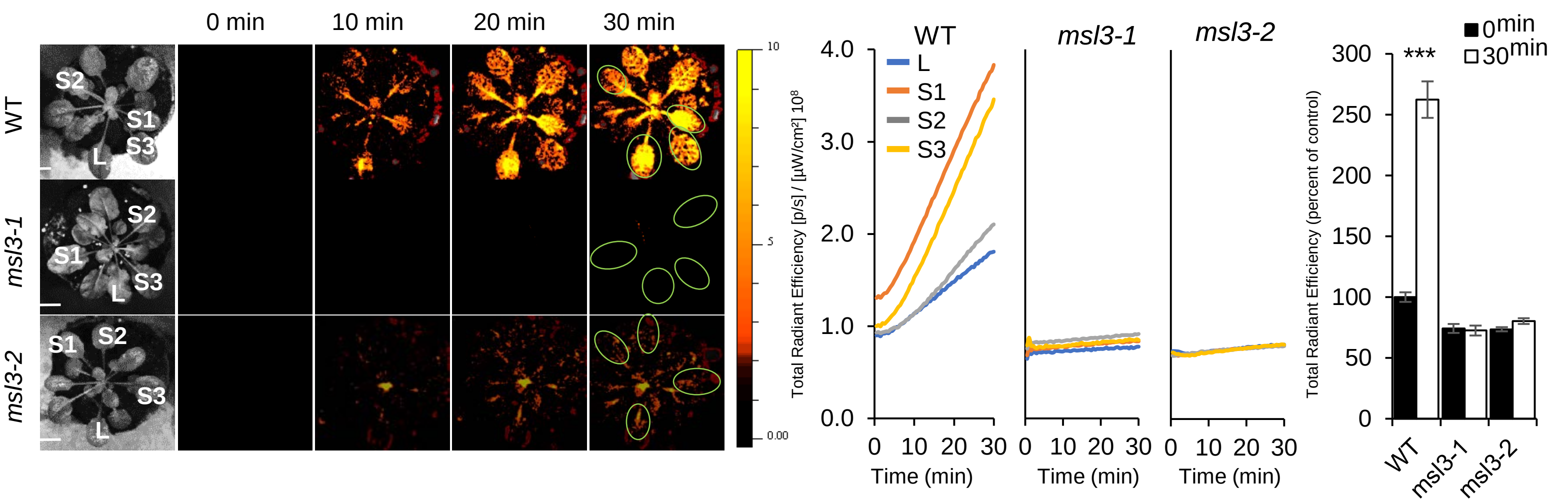


**Fig. S2. The calcium-permeable channel MSL3 is required for mediating light stress-induced systemic signaling in Arabidopsis.**

Representative time-lapse images of systemic ROS accumulation in wild-type and *msl3* (two independent alleles) *Arabidopsis thaliana* plants subjected to a 2 min local (L) high light (HL) stress treatment (applied to leaf L only), is shown on left; representative continuous measurements of ROS levels in the local (L) and systemic (S) leaves over the entire course of the experiment are shown in the middle (ROIs used for calculating them are indicated with light green circles on the images to the left); and statistical analysis of ROS accumulation in local and systemic leaves of all plants used for the analysis at 0 and 30 min is shown on right. All experiments were repeated at least 3 times with 10 plants per biological repeat. Student t-test, SE, N=12, ***P < 0.005. Scale bar indicates 1 cm. *Abbreviations used:* HL, high light; L, local; MSL, mechanosensitive channel of small conductance-like; ROI, region of interest; S, systemic.

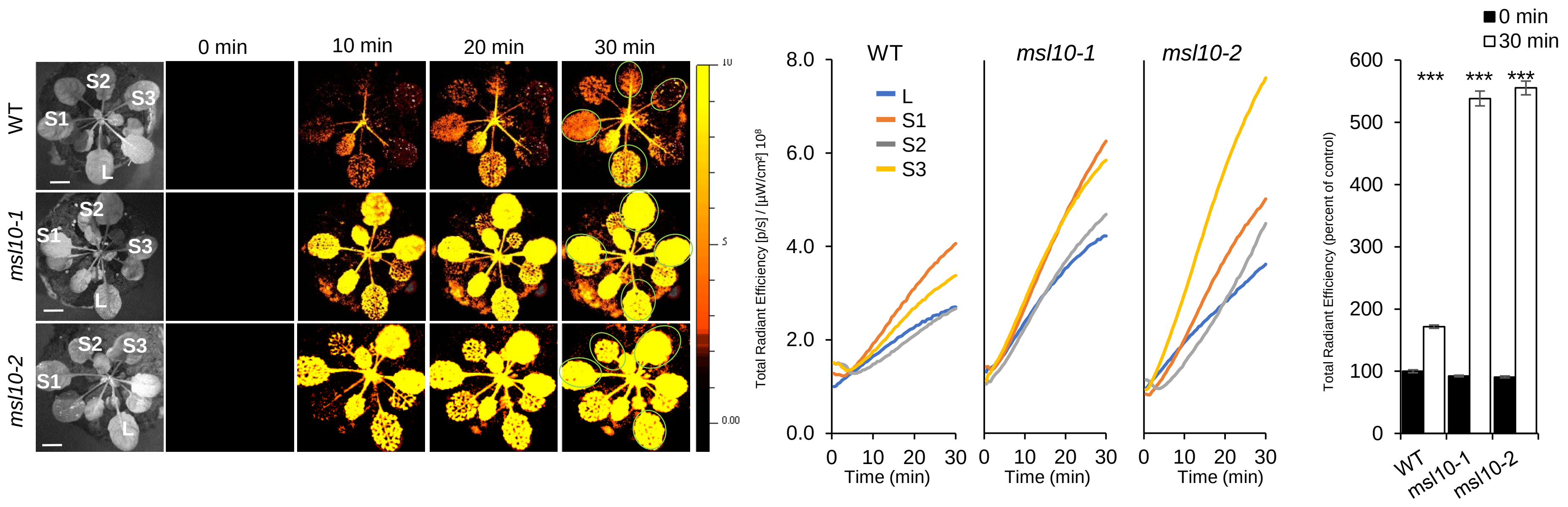


**Fig. S3. Enhanced systemic ROS signal in mutants impaired in the MSL10 calcium-permeable channel during light stress-induced systemic signaling in Arabidopsis.**

Time-lapse imaging of systemic ROS accumulation in wild-type and *msl10* (two independent alleles) *Arabidopsis thaliana* plants subjected to a 2 min local (L) high light (HL) stress treatment (applied to leaf L only), is shown on left; continuous measurements of ROS levels in the local (L) and systemic (S) leaves over the entire course of the experiment are shown in the middle (ROIs used for calculating them are indicated with light green circles on the images to the left); and statistical analysis of ROS accumulation in local and systemic leaves of all plants used for the analysis at 0 and 30 min is shown on right. All experiments were repeated at least 3 times with 10 plants per biological repeat. Student t-test, SE, N=12, ***P < 0.005. Scale bar indicates 1 cm. *Abbreviations used:* HL, high light; L, local; MSL, mechanosensitive channel of small conductance-like; ROI, region of interest; S, systemic.

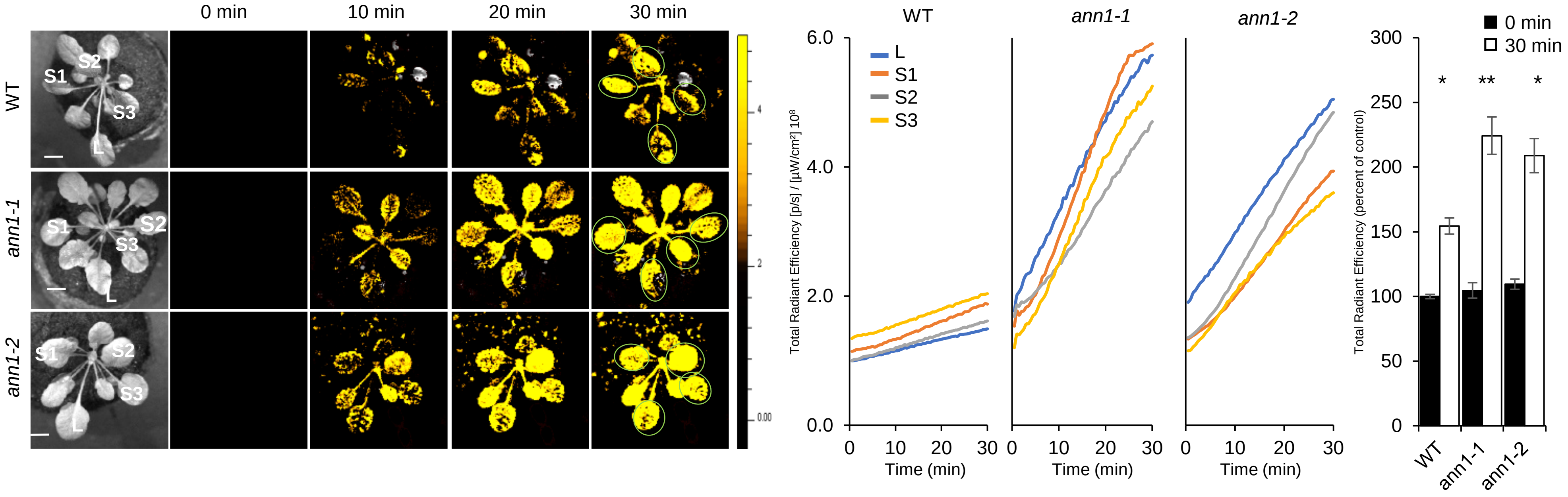


**Fig. S4. Enhanced systemic ROS signal in mutants impaired in the ANN1 calcium-permeable channel during light stress-induced systemic signaling in Arabidopsis.**

Representative Time-lapse images of systemic ROS accumulation in wild-type and *ann1* (two independent alleles) *Arabidopsis thaliana* plants subjected to a 2 min local (L) high light (HL) stress treatment (applied to leaf L only), is shown on left; representatives line graphs of continuous measurements of ROS levels in the local (L) and systemic (S) leaves over the entire course of the experiment are shown in the middle (ROIs used for calculating them are indicated with light green circles on the images to the left); and statistical analysis of ROS accumulation in local and systemic leaves of all plants used for the analysis at 0 and 30 min is shown on right. All experiments were repeated at least 3 times with 10 plants per biological repeat. Student t-test, SE, N=12, $^*P < 0.05$, $^{**}P < 0.01$. Scale bar indicates 1 cm. *Abbreviations used:* HL, high light; L, local; ANN, annexin; ROI, region of interest; S, systemic.

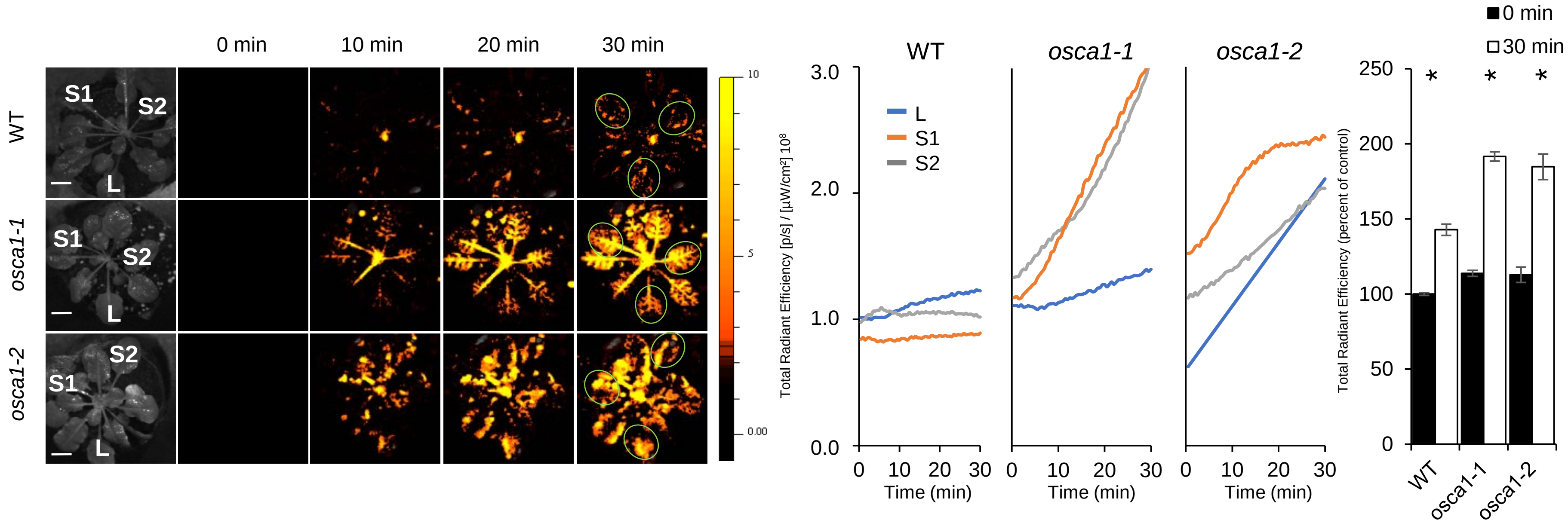


**Fig. S5: Enhanced systemic ROS signal in mutants impaired in the OSCA1 calcium-permeable channel during light stress-induced systemic signaling in Arabidopsis.**

Time-lapse imaging of systemic ROS accumulation in wild-type and *gat1* (two independent alleles) *Arabidopsis thaliana* plants subjected to a 2 min local (L) high light (HL) stress treatment (applied to leaf L only), is shown on left; continuous measurements of ROS levels in the local (L) and systemic (S) leaves over the entire course of the experiment are shown in the middle (ROIs used for calculating them are indicated with light green circles on the images to the left); and statistical analysis of ROS accumulation in local and systemic leaves of all plants used for the analysis at 0 and 30 min is shown on right. All experiments were repeated at least 3 times with 10 plants per biological repeat. Student t-test, SE, N=12, *P < 0.05. Scale bar indicates 1 cm. *Abbreviations used:* HL, high light; L, local; OSCA, reduced hyperosmolality-induced [Ca(2+)]i increase 1; ROI, region of interest; S, systemic.

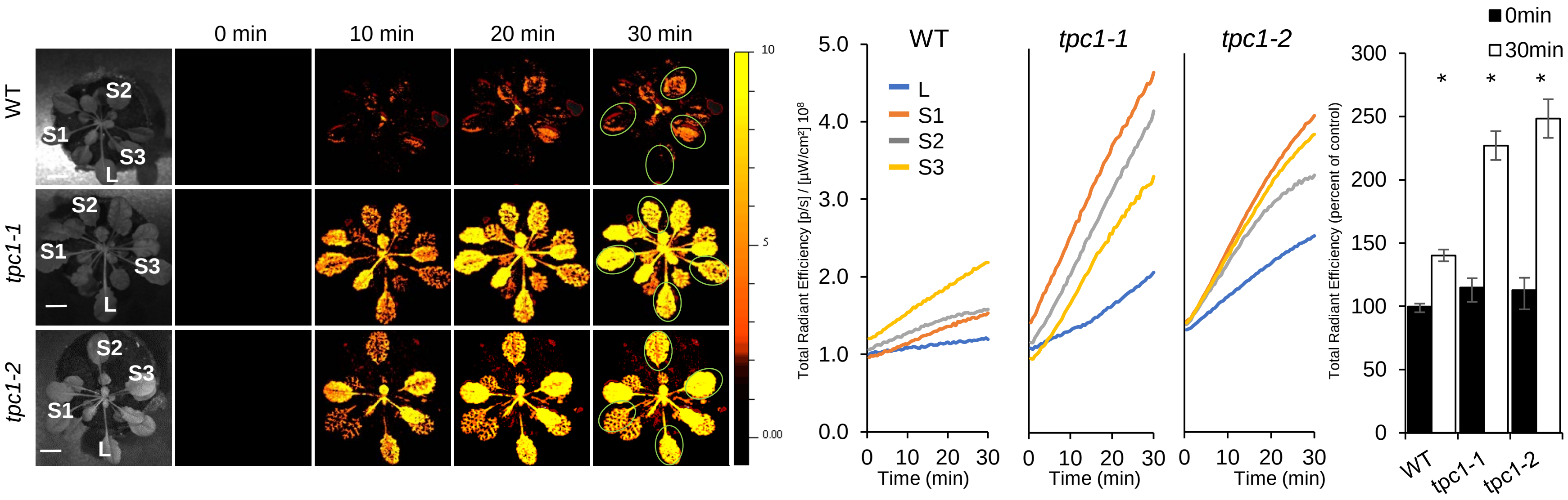


**Fig. S6. Enhanced systemic ROS signal in mutants impaired in the TPC1 calcium-permeable channel during light stress-induced systemic signaling in Arabidopsis.**

Time-lapse imaging of systemic ROS accumulation in wild-type and *tpc1* (two independent alleles) *Arabidopsis thaliana* plants subjected to a 2 min local (L) high light (HL) stress treatment (applied to leaf L only), is shown on left; continuous measurements of ROS levels in the local (L) and systemic (S) leaves over the entire course of the experiment are shown in the middle (ROIs used for calculating them are indicated with light green circles on the images to the left); and statistical analysis of ROS accumulation in local and systemic leaves of all plants used for the analysis at 0 and 30 min is shown on right. All experiments were repeated at least 3 times with 10 plants per biological repeat. Student t-test, SE, N=12, *P < 0.05. Scale bar indicates 1 cm. *Abbreviations used:* HL, high light; L, local; TPC, two-pore channel; ROI, region of interest; S, systemic.

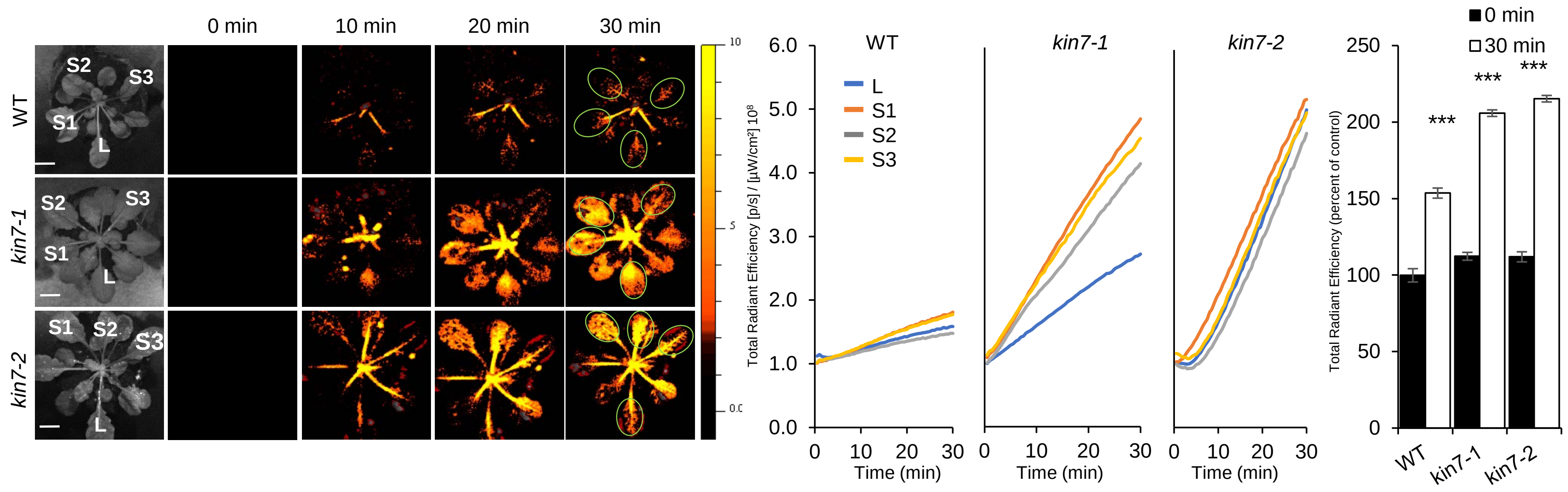


**Fig. S7. Enhanced systemic ROS signal in mutants impaired in the plasmodesmata-regulating protein KIN7 during light stress-induced systemic signaling in Arabidopsis.**

Time-lapse imaging of systemic ROS accumulation in wild-type and *kin7* (two independent alleles) *Arabidopsis thaliana* plants subjected to a 2 min local (L) high light (HL) stress treatment (applied to leaf L only), is shown on left; continuous measurements of ROS levels in the local (L) and systemic (S) leaves over the entire course of the experiment are shown in the middle (ROIs used for calculating them are indicated with light green circles on the images to the left); and statistical analysis of ROS accumulation in local and systemic leaves of all plants used for the analysis at 0 and 30 min is shown on right. All experiments were repeated at least 3 times with 10 plants per biological repeat. Student t-test, SE, N=12, ***$P < 0.005$. Scale bar indicates 1 cm. *Abbreviations used:* HL, high light; L, local; KIN7, kinase 7; ROI, region of interest; S, systemic.

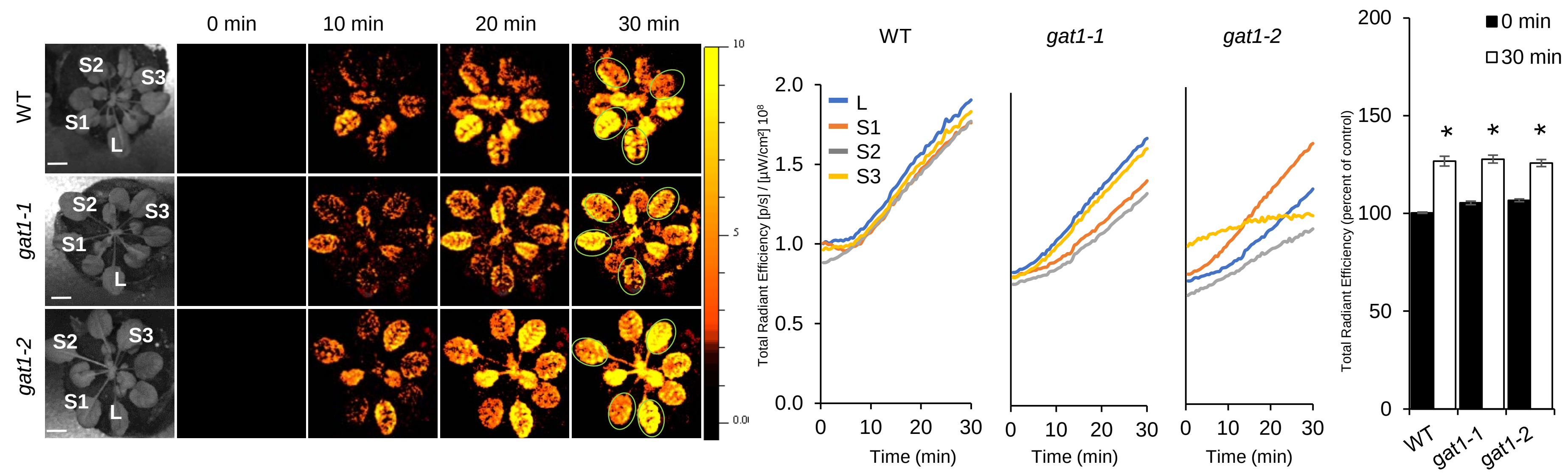


**Fig. S8. Plasmodesmata-regulating protein GAT1 is not required for mediating light stress-induced systemic signaling in Arabidopsis.**

Time-lapse imaging of systemic ROS accumulation in wild-type and *gat1* (two independent alleles) *Arabidopsis thaliana* plants subjected to a 2 min local (L) high light (HL) stress treatment (applied to leaf L only), is shown on left; continuous measurements of ROS levels in the local (L) and systemic (S) leaves over the entire course of the experiment are shown in the middle (ROIs used for calculating them are indicated with light green circles on the images to the left); and statistical analysis of ROS accumulation in local and systemic leaves of all plants used for the analysis at 0 and 30 min is shown on right. All experiments were repeated at least 3 times with 10 plants per biological repeat. Student t-test, SE, N=12, *P < 0.05. Scale bar indicates 1 cm. *Abbreviations used:* HL, high light; L, local; GAT, gfp arrested trafficking; ROI, region of interest; S, systemic.

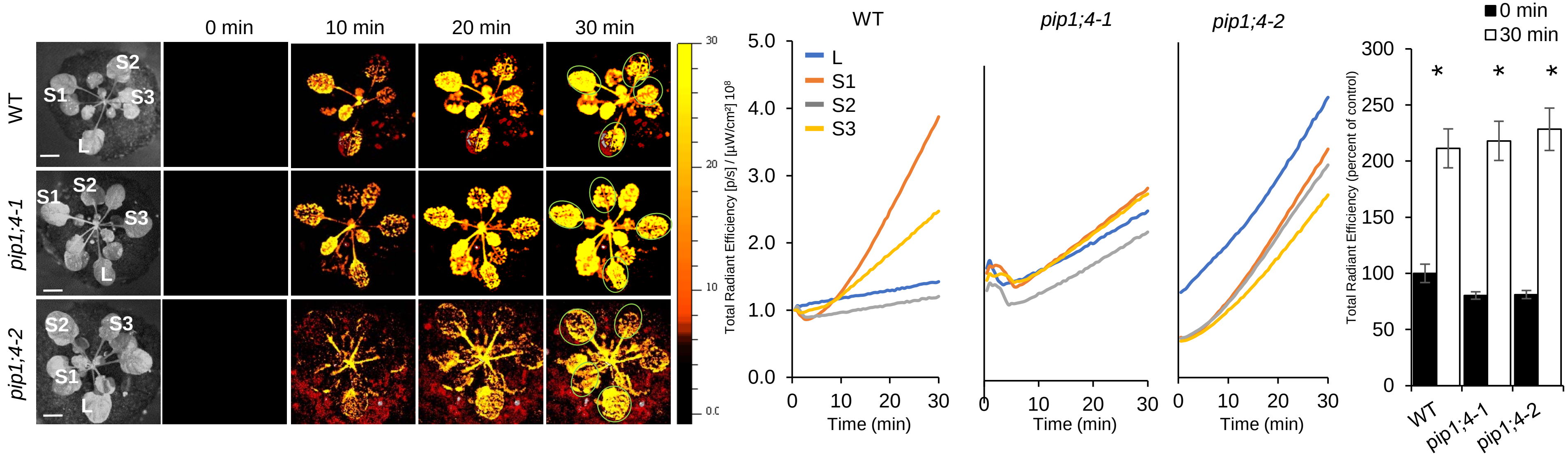


**Fig. S9. Aquaporin PIP1;4 is not required for mediating light stress-induced systemic signaling in Arabidopsis.**

Time-lapse imaging of systemic ROS accumulation in wild-type and *pip1;4* (two independent alleles) *Arabidopsis thaliana* plants subjected to a 2 min local (L) high light (HL) stress treatment (applied to leaf L only), is shown on left; continuous measurements of ROS levels in the local (L) and systemic (S) leaves over the entire course of the experiment are shown in the middle (ROIs used for calculating them are indicated with light green circles on the images to the left); and statistical analysis of ROS accumulation in local and systemic leaves of all plants used for the analysis at 0 and 30 min is shown on right. All experiments were repeated at least 3 times with 10 plants per biological repeat. Student t-test, SE, N=12, *P < 0.05. Scale bar indicates 1 cm. *Abbreviations used:* HL, high light; L, local; PIP, plasma membrane intrinsic protein; ROI, region of interest; S, systemic.

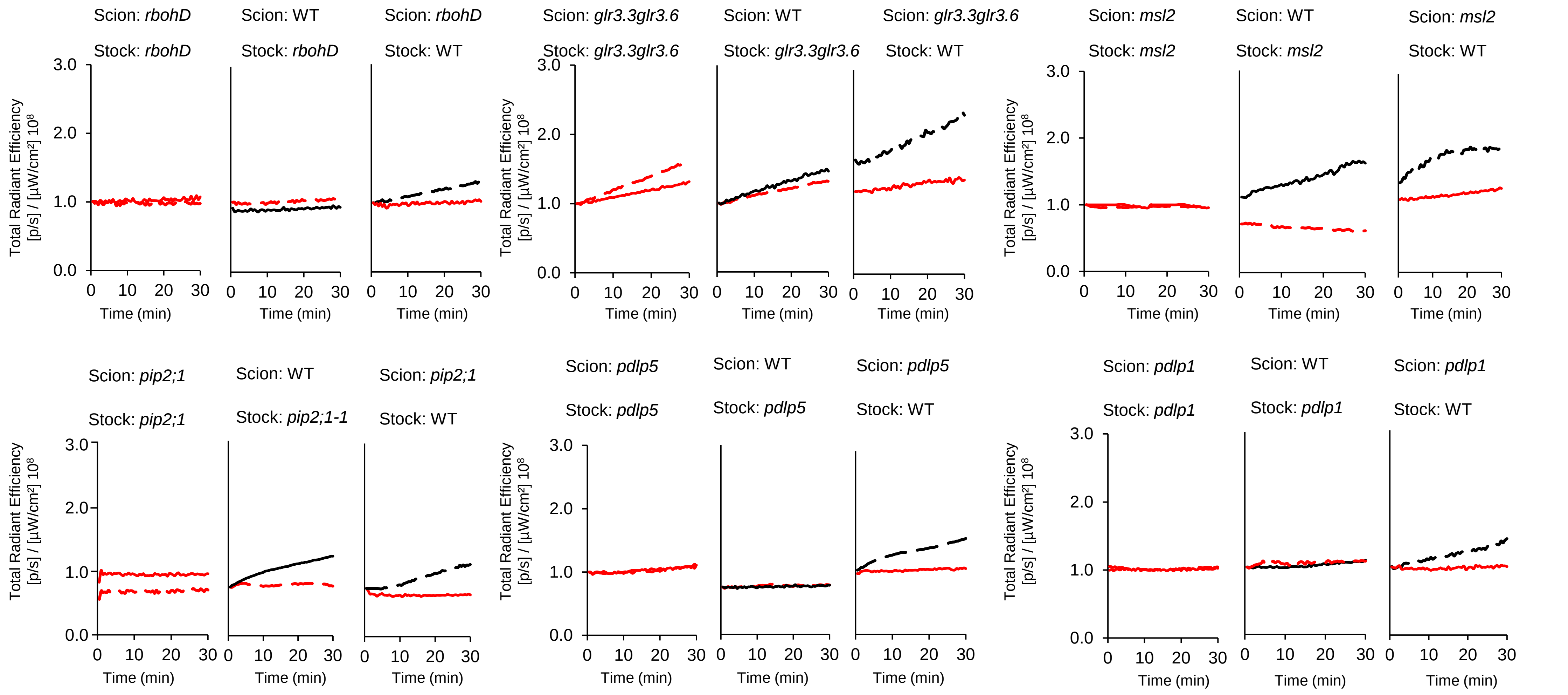


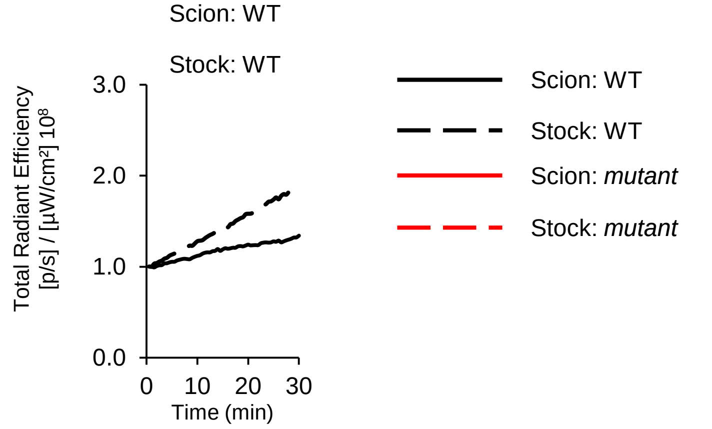


**Fig. S10: Plasmodesmata function and apoplastic ROS production are required for both initiation and propagation of the systemic signal.**

Representative ROS signal intensity line graphs in the stock and scion of different grating combinations between wild-type and mutants impaired in systemic signaling (*rbohD*, *glr3.3glr3.6*, *cngc2*, *msl2*, *pip2;1 pdlp5* and *pdlp1*), following a 2 min high light stress treatment of a single stock leaf (local leaf). *Abbreviations* used: CNGC, cyclic nucleotide-gated ion channel; GLR, glutamate receptor-like; MSL, mechanosensitive channel of small conductance-like; PDLP, plasmodesmata localized protein; PIP, plasma membrane intrinsic protein; RBOHD, respiratory oxidase burst homolog D; ROS, reactive oxygen species; Sc, scion; St. Stock; WT, wild-type.

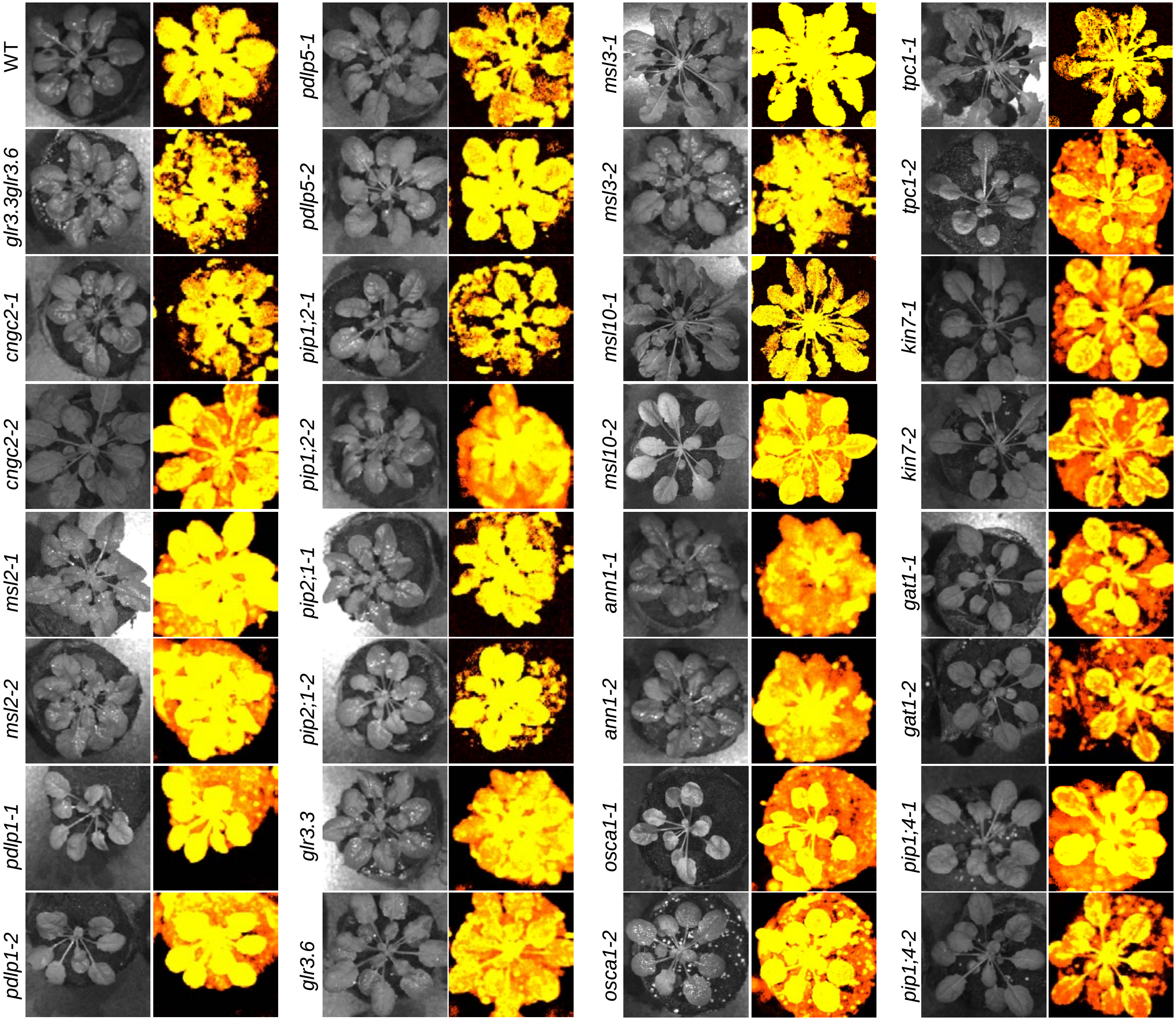


**Fig. S11: Systemic ROS imaging following hydrogen peroxide fumigation in the tested plants (as control for dye penetration in the different mutants).**

Plants were fumigated with DCF according to *(8)*, and then fumigated for 10 min with 0.3% $H_2O_2$. Images were acquired with IVIS Lumina S5 and analyzed with Living Image software.
